# The human default consciousness and its disruption: insights from an EEG study of Buddhist jhāna meditation

**DOI:** 10.1101/407544

**Authors:** Paul Dennison

**Affiliations:** Paul Dennison Psychotherapy

**Keywords:** EEG, meditation, jhāna, consciousness, epilepsy, infraslow-waves, spike-waves, free-energy, active inference

## Abstract

The “neural correlates of consciousness” (NCC) is a familiar topic in neuroscience, overlapping with research on the brain’s “default mode network”. Task-based studies of NCC by their nature recruit one part of the cortical network to study another, and are therefore both limited and compromised in what they can reveal about consciousness itself. The form of consciousness explored in such research, we term the human default consciousness (DCs), our everyday waking consciousness. In contrast, studies of anaesthesia, coma, deep sleep, or some extreme pathological states such as epilepsy, reveal very different cortical activity; all of which states are essentially involuntary, and generally regarded as “unconscious”. An exception to involuntary disruption of consciousness is Buddhist jhāna meditation, whose implicit aim is to intentionally withdraw from the default consciousness, to an inward-directed state of stillness referred to as jhāna consciousness, as a basis to develop insight. The default consciousness is sensorily-based, where information about, and our experience of, the outer world is evaluated against personal and organic needs and forms the basis of our ongoing self-experience. This view conforms both to Buddhist models, and to the emerging work on active inference and minimisation of free energy in determining the network balance of the human default consciousness.

This paper is a preliminary report on the first detailed EEG study of jhāna meditation, with findings radically different to studies of more familiar, less focused forms of meditation. While remaining highly alert and “present” in their subjective experience, a high proportion of subjects display “spindle” activity in their EEG, superficially similar to sleep spindles of stage 2 nREM sleep, while more-experienced subjects display high voltage infraslow-waves reminiscent, but significantly different, to the slow waves of deeper stage 4 nREM sleep, or even high-voltage delta coma. Some others show brief posterior spike-wave bursts, again similar, but with significant differences, to absence epilepsy. Some subjects also develop the ability to consciously evoke clonic seizure-like activity at will, under full control. We suggest that the remarkable nature of these observations reflects a profound disruption of the human DCs when the personal element is progressively withdrawn.

## 1. INTRODUCTION

### 1.1 A DEFAULT SENSORY CONSCIOUSNESS (DCs)

We consider the DCs to be a sensory consciousness, irrespective of whether we are responding to direct sensory input, accessing memory or processing ideas, or indeed dreaming, all of which are experienced within a sensory framework. This consciousness is represented at the cortical and biological level as an ongoing dynamic functional organisation intrinsic to our lives as human beings interacting with others and the world. This default consciousness is the *de facto* subject of the many “content” studies of neural correlates of consciousness (NCC) where researchers examine which cortical networks are stimulated or suppressed by a subject undergoing tasks or external stimuli (Koch *et al*., 2016; Boly *et al*., 2017).

A subjective component is central to the DCs, and is likely the dominant factor in the dynamic neuronal balance between inputs from the outer world and from our body, with their resonances to past experiences held in memory, then weighed as to their value to the “I” or “self” in the light of current needs or actions. This understanding of consciousness is implicit in psychoanalysis, from Freud’s early work in his *Project for a Scientific Psychology* (1895), to the clinical experience of psychotherapists and psychoanalysts of the constant ongoing resonance between current and past experience held in memory, memories of reciprocal roles that contain information on the emotional impact of events (pain-pleasure; liking-disliking) for the self (e.g. Ryle and Kerr, 2002; and Matte Blanco’s, 1980, description of the *Unconscious as Infinite Sets*). It also conforms to the emerging work within neuroscience on active inference, interoceptive inference and selfhood (Friston *et al*., 2016; Seth *et al*., 2012, 2016).

Whilst content studies of the NCC can reveal different features of the DCs, state-based approaches compare it to states such as sleep, anaesthesia, coma, epilepsy and some pathological states, mostly regarded as unconscious (Boly *et al*., 2011; Gosseries *et al*., 2014; Owen *et al*., 2006). If it is possible, as we aim to demonstrate, for a person to intentionally and progressively withdraw their personal involvement from the DCs, even partially, while remaining fully conscious, a new window is opened into exploring the NCC (Hohwy, 2009), and consciousness itself.

### 1.2 JHĀNA MEDITATION

Given the rather remarkable observations we describe, certainly atypical compared to previous EEG studies of meditation, it is appropriate to give a context and overview of some of the key features of Buddhist jhāna meditation.

Whether Southeast Asian, Tibetan, Japanese or Chinese, Buddhist meditation comprises two strands, Samatha (Wallace, 1999) and Vipassanā (Cousins, 1994-96); the former often translated as tranquillity or serenity, and the latter as insight or wisdom. Mindfulness, though well-known as a form of practice in its own right, and accepted as useful in the treatment of recurrent depression, is just one of the basic factors underpinning both samatha and vipassanā. Jhāna meditation falls within the samatha division. Etymologically, jhāna is often translated as “absorption”, but has a secondary root, *jhapeti*, meaning to burn up, which is a reflection that it is a highly active and energised state (Cousins, 1973; Gunaratana, 1980). While there have been many EEG studies of meditation (Thomas and Cohen, 2014; Cahn and Polich, 2006), there have been no in-depth studies of jhāna meditation, and there exist very few centres that teach its practice. The background to this requires some explanation. In South and Southeast Asia during the 1950s, Buddhist practices underwent a “Reform”, where age-old samatha practices were criticised as unscientific, and repressed in favour of a heavily politically-promoted form of meditation known as Burmese vipassanā, which claimed that jhāna was not necessary as a prerequisite for insight and realisation of Buddhist goals (Crosby, 2013). As a result, jhāna meditation was relegated to an almost esoteric role, sidelined in favour of vipassanā. Many also believed it was not possible to develop and practice jhāna in a lay context, outside monastic and forest meditation traditions.

Recently, an interest in jhāna has revived, with two main traditions emerging in the West. The first, and best known, comes via monastic teachers, and in this form the breath is not controlled in a formal manner. Hagerty *et al.* (2013) describe an EEG study of a single Western practitioner of this method, but with results very different to those we find. In the second form, the length of breath *is* controlled in the approach to jhāna, since the “normal” length is regarded as integral to the DCs, withdrawal from which is the primary aim of jhāna. This form was introduced to the UK in the 1960s by a former Thai-Cambodian Buddhist monk (The Samatha Trust, UK regd Charity 1973), and is closely related to the *Yogāvacara*, a formerly widespread and ancient, mainly oral, tradition of meditation, practiced by both monks and lay people across South and Southeast Asia, currently the subject of research by ethnologists based on palm-leaf manuscripts discovered in Cambodia and Thailand (Bizot, 1994; Crosby, 2000). In some monasteries, *Yogāvacara* techniques were regarded as a means to develop mastery of jhāna, but were also considered particularly suitable for lay meditators leading normal household lives, as are the subjects of this study. Using lengths of breath longer or shorter than “normal”, marks a protected space where jhāna can be safely developed, and safely left to return to the normal DCs and daily life without conflict. “Safely”, refers to the containment of highly energised states frequently developed in approaching jhāna; in this respect, and in the precise ways in which the breath is controlled, there are similarities to Tibetan Buddhist yoga (Minvaleev, 2014).

Wallace (1999), referring to the samatha tradition and the practice of jhāna, comments that “The mind and consciousness itself are the primary subjects of introspective investigation within the Buddhist tradition”. Indeed, the techniques employed are remarkably relevant to current neuroscience, once the different terminologies are understood. To develop jhāna, the meditator’s first task is to overcome the “hindrances”, usually listed as: sense-desire, ill-will, sloth and torpor, restlessness and worry, and doubt (Cousins, 1973; Gunaratana, 1980). These represent, in our terms, key features of the human DCs: a constant evaluation of sensory input against past experience and future needs and expectations, based on value to the “I” or “self”, which implies liking or disliking, pleasure versus pain, habits of attachment, as well as available energy, restlessness and doubt. In practice, a meditator does not think about or dwell on the hindrances during meditation, but simply begins by turning attention to the breath; each time the mind wanders, distractions in the form of thoughts or feelings are acknowledged and attention patiently, again and again, brought back to the breath. The Buddhist jhāna tradition describes eight jhānas: four rūpa (form) jhānas, and four arūpa (formless) jhānas; of which this paper deals with the former.

#### 1.2.1. Attention

The jhānas are described by their characterising factors; 5 factors for the first rūpa jhāna, 3 for the second, and 2 for the third and fourth jhānas, as listed below with the Pali terms:

First rūpa jhāna factors

- <u>Applied</u> attention, or initial thought (= *vitakka*)
- <u>Sustained</u> attention or thought (= *vicāra*)
- Energised interest, or “joy” (= *pīti*)
- Happiness, contentment or bliss (= *sukha*)
- One-pointedness of mind (= *ekaggatācitta*)

Second rūpa jhāna factors

- Energised interest, or “joy” (= *pīti*)
- Happiness, contentment or bliss (= *sukha*)
- One-pointedness of mind (= *ekaggatācitta*)

Third rūpa jhāna factors

- Happiness, contentment or bliss (= *sukha*)
- One-pointedness of mind (= *ekaggatācitta*)

Fourth rūpa jhāna factors

- One-pointedness of mind (= *ekaggatācitta*)
- Equanimity(= *upekkha*)

The jhānas are a progressive sequence towards deeper states of equanimity, or serenity. The dominant factors of the first rūpa jhāna are two aspects of attention: *vitakka*, applied attention, is the repeated placing of attention on the meditation object, in this case the breath; and *vicāra*, sustained attention, develops as the meditator becomes more skilled at noticing and resisting distraction. Once attention is stabilised internally, the meditator can progress towards the second, third and fourth jhānas, which are more feeling-based, and share an increasing sense of equanimity. Working with attention is the dominant activity in developing the first jhāna, and is required to be developed to a high level, often over years of practice. Typically, practitioners start by mentally counting during in and out breaths, to aid noticing when attention wanders. Distractions are acknowledged minimally, before returning to the count. Maintaining different lengths of breath aids mindfulness, and four lengths are used, two longer than “normal”, and two shorter. As distraction eases, counting is dropped, replaced by following the breath from its touch at the nose tip, through the sensations at the throat, chest and diaphragm, and back on the out-breath; noting and managing distractions. This phase is described as developing access concentration. Finally the meditator rests attention at one point, usually the nose tip, and at this stage attention is progressively transferred to an internal mental object, referred to in the jhāna texts as the *nimitta* (Wallace, 1999; Cousins, 1973) or “sign”, which is the meditator’s growing sense of his/her own consciousness. We are tempted to say the “qualia” of that consciousness, provided qualia is not interpreted in sensory terms as too-crudely “this” or “that”.

Allowing for terminology, we expect these two factors of attention to have counterparts in the executive attention networks of the brain. However, attention is inseparable from broader processes of perception (Hohwy 2012), which requires us to consider the personal component, and the subjective experience of the meditator is a clue as to what we might expect. At first a meditator’s attention, as in the DCs, is strongly sensorily-determined by the habit or need to mentally “commentate”, “name” or “recognise”, i.e. to orient experience within the familiar sensory structure of the DCs. Since subjectivity in these perceptual processes is heavily “Eye/I”-driven, we may expect disruption to the executive attention networks, but also to the ventral and dorsal perceptual streams (Milner, 2017; Cloutman, 2012), as the meditator resists the pull back towards DCs processes in developing the first rūpa jhāna.

#### 1.2.2. Attachment

Development of the second, third and fourth rūpa jhānas is more concerned with feeling, and the underlying subject-object nature of consciousness rather than the cognitive processes of attention. In fact, even to develop the first rūpa jhāna, meditators are already working with resisting attachment to liking and disliking, which in Buddhist terms are the roots of craving and the source of suffering. Here there is a correspondence to Freud’s “pleasure principle”, and the twin pulls of craving and aversion, as well as the importance of understanding attachment disorders in psychiatry and psychotherapy. Since liking and disliking, and associated emotions, are dominant features of our habitual DCs, linking perception to action choices, it may be no surprise to find rather dramatic changes in brain activity when the personal element is withdrawn from the DCs.

Subjectively, the movement from the first to the second rūpa jhāna is characterised by a growing sense of peace and contentment, at the increasing freedom from dependence on DCs processes, as well as a growing “energised interest”. These are the two factors *sukha* and *pīti*, respectively, listed above for the second rūpa jhāna, together with the third factor one-pointedness of mind (*ekaggatācitta*) which underpins all four jhānas. The factor *pīti* is strongly emphasised in the Yogāvacara, which lists 5 levels of intensity ranging from fine bodily vibration or prickling of the hairs on the head and body, to, at its most intense, bodily shaking and even jumping in the air (*The Yogāvachara’s Manual*). *Pīti* represents the growing involvement of the body and subtle bodily energies into a state of mind-body integration referred to as *samādhi*, a term frequently used interchangeably with “concentration” or jhāna. The EEG observations of high energy states described in this paper, are, we believe, the counterparts of this energisation, and signal the beginnings of a transition to the second and higher rūpa jhānas.

#### 1.2.3. Subject-Object

Whilst the task of the first rūpa jhāna is to develop a degree of mastery of attention, the task of the second rūpa jhāna is to master *pīti*; not to suppress it, but to tranquilise (Pali, *passaddhi*) any bodily disturbance into an increasingly still mental state “held” by attention to the *nimitta*. For some meditators the *nimitta* is experienced visually as light, for others as touching something intangible with the mind, or by others as listening to silence (Buddhaghosa, *Visuddhimagga*); these differing experiences reflecting the traces of a meditator’s habitual preference for one or other sense modality in their DCs. Once *pīti* is tranquilised and incorporated, the third rūpa jhāna can develop, and is described in 5^th^-century texts as being “completely conscious” (Upatissa, *Vimuttimagga*), or with “full awareness like that of a man on a razor’s edge” (Buddhaghosa, *Visuddhimagga*). Interestingly, avoiding the words conscious or aware, subjects of this study prefer, “presence”, or “vivid presence” for their subjective experience.

Practice of the second, third and fourth rūpa jhānas is also regarded as a progressive exploration and refinement of the subject-object relationship (*nāma-rūpa*, or name and form in Buddhist terms), which becomes less cognitively and sensorily-determined, and less dependent on liking or disliking; such that in the fourth rūpa jhāna even dependence on the “reward” of pleasure or satisfaction ends, replaced by deep stillness and finely poised balance and equanimity. The nature of the subject-object experience is of considerable interest to neuroscience, and will be taken up later in this paper.

However, to develop the jhāna factors is not a straightforward cognitive process, as in task-based EEG studies; meditators cannot simply “think” themselves into jhāna. While the motivation is to withdraw from the habitual DCs (“*secluded from sense desire*…”, Cousins, 1973), it is the *nimitta* acting as an “attractor” that allows meditators to settle into jhāna. It could be said that the *nimitta* plays a similar role to the feedback sign in neurofeedback (Sitaram R. *et al*., 2017), and that Samatha meditation is a naturalistic form of neurofeedback, predating modern forms by over two millennia (Dennison, 2012).

## 2. METHODS AND MATERIALS

### 2.1. SUBJECTS AND RECORDING PROTOCOL

It is important to acknowledge that meditators in this study, while very experienced in samatha meditation, vary considerably in their experience of jhāna. Many see jhāna as a progressive process to develop a solid base of equanimity prior to developing insight (vipassanā), and not all are overly concerned to develop mastery of the subtleties between the different jhānas, while leading busy day-to-day lives. In fact none of our subjects would claim complete mastery of jhāna, not surprising given that such mastery is regarded as a rare achievement in Buddhism, described in the 5^th^ century *Visuddhimagga* as of three levels, inferior, medium and superior. In a pilot study in 2010-14, we did not find unequivocal EEG signatures of the different jhānas based on subjective feedback, despite finding startling EEG activity suggesting this form of meditation has profound effects on brain activity.

Accordingly, in this paper we choose not to rely <u>primarily</u> on subjective views as to which jhāna meditators feel they are experiencing, but focus rather on the <u>themes</u> of EEG activity that emerge across this broad group of meditators as they attempt to develop ever-deeper stages of meditation. In effect, the material constitutes a series of case studies describing paroxysmal electrophysiological changes in the EEG; i.e., spindles, infraslow waves, spike-wave bursts and clonic seizure-like activity, never previously observed in other researched forms of meditation. The Results section will explore these features in detail in both sensor space and source space. Since the common factor underlying these themes is the practice of jhāna meditation, we hold in mind a hypothesis that these effects may be related to the nature of jhāna, in particular to the underlying goal of jhāna meditation to withdraw the personal component from our default sensory consciousness. We therefore take a cross-discipline approach to explore this hypothesis by comparing the EEG evidence to detailed Buddhist understandings of the characteristics of the different rūpa jhānas in the Discussion section.

Epochs of these paroxysmal features were observed in every subject to a greater or lesser degree, and we explore the neural correlates of each of these themes in both sensor space and source space (following independent component analysis and source reconstruction using eLoreta). We adopt a common protocol for all meditators, using verbal cues to first record a few minutes of resting state eyes-closed and eyes-open EEG, followed by meditators attempting to progressively develop the jhānas, for a total recording time of 35-40 mins. For some meditators this protocol is much quicker than their everyday practice, with corresponding disadvantages, but is adopted for consistency. Subjects practice seated on the ground, usually on a cushion, with the body unsupported, erect and composed. The observer/researcher notes events such as shifts in posture, a cough, external noise, or anything likely to cause an artifact.

This is a within-group study, where the control group is, in effect, the wealth of other EEG studies of meditation and the NCC. Twenty-nine experienced meditators (19 men, 10 women) from a total pool of around 400 in the UK, Ireland and the US, all experienced in samatha meditation as taught by the Samatha Trust, a UK registered charitable organisation, were recorded during 2014-18, some also re-recorded after 1-3 year intervals. Years’ experience of samatha meditation range from 4–40+ years, with most individuals maintaining a daily practise of 30-60 mins, with more intensive 7-10 days “retreats” every 1–3 years. Twenty-four of the subjects are of graduate or postgraduate-levels of education, and more than half hold, or have held, senior professional roles in health-care or education. Four have spent temporary periods from 1-10 months as ordained Buddhist monks in Thailand. As noted earlier, while all subjects are very experienced in samatha meditation, experience of jhāna varies considerably.

Given that recordings show features superficially similar to unconscious states, we stress that subjects are fully conscious throughout, with no signs of loss of muscle tone or sleepiness in posture. Subjects respond quickly to verbal cues; and finally show no signs of sleep inertia or disorientation. In fact, meditators describe feeling more alert and present during and after practice. Following a recording subjects recollect their practice while the researcher monitors the recording in parallel.

### 2.2. EQUIPMENT AND ANALYSIS

Recordings were made using 24-bit Mitsar DC amplifiers, either the 31-channel Mitsar 202, or the 21-channel wireless SmartBCI amplifier (Mitsar Medical, St Petersburg). The former having a sampling rate of 500/sec and upper frequency limit of 150 Hz, used with Easycaps and sintered Ag/AgCl electrodes; the latter a sampling rate of 250/sec and upper frequency limit 70 Hz, used with MCScaps and sintered Ag/AgCl electrodes. In using DC amplifiers, choosing the best cap/gel combination is critical to minimise drift and maintain stable low impedances at the electrode-skin interface. Having tested many combinations, we agree with Tallgren *et al*. (2005) that saline-based gels used with sintered Ag/AgCl electrodes are essential with DC amplifiers, and after testing most commercial gels, we favour a combined conductive and mildly abrasive gel to obtain impedances close to or less than 5KΩ, with caps that allow good visual access to the electrode-skin interface to apply gel.

Electrodes were placed according to the international 10-20 system, with a monopolar linked-ears reference. Software analysis was carried out using WinEEG, implementing the Infomax algorithm as used in EEGLAB (Delorme and Makeig, 2004) to compute spectra and independent components (ICs). Cortical sources were computed using the reverse solution of eLoreta (Pascual-Marqui, 2007). Bandwidth and epoch length are important variables in computing spectra and ICs for the different features we observe; that is, spindles, infraslow waves (ISWs) and spike-waves. Since the ISWs we analyse are very strong against the EEG background, we adopt a broad bandwidth of 0.032-70/150 Hz, depending on the amplifier model, to capture the full range of spectral activity; whereas for spindles a bandwidth of 5.3-15 Hz is used to minimise confusion with ISWs and higher frequency beta and gamma activity; and for spike waves, 0.53-70/150 Hz, again to minimise ISW confusion, but to retain high frequency content given the harmonic structure of the spike waves.

Epoch length introduces a smoothing effect of its own irrespective of the pass-band, such that epochs of 4, 8, 16, 32 and 64 secs smooth frequencies below 0.25, 0.125, 0.063, 0.031 and 0.016 Hz respectively. To capture ISW activity, we use epochs of 16 or 32 secs depending on the length of segment analysed; for spindles we use 4 secs; and for the very brief spike-wave bursts we use the shortest WinEEG epoch of 1 sec; all with 50% overlap Hanning windows.

### 2.3. ARTIFACTS OR CORTICAL?

With atypical EEG phenomena, the question of what is artifact and what is cortical is very important, with the risk of discarding important cortical activity if some unusual feature is too quickly labelled “artifact”. Accordingly, the experimental design aims, as far as possible, to prevent artifacts arising in the first place. In an early pilot study, movement and eye artifacts were a concern, the former caused mainly by movement of electrode connector wires if a subject moves during recording. For the 31-electrode system the most effective solution has been to gather electrode wires into an “umbilical cord” sheathed in earthed carbon-fibre material between headcap and amplifier; whereas the 21-channel system is already resilient to movement artifacts due to short connections to the wireless amplifier mounted directly on the headcap. Combined with subjects experienced in meditation, used to holding still postures for long periods, as well as the observer/researcher noting posture changes or signs of muscle tension, very few segments were excluded. The only situation where movement remains a problem is when recording the deliberate arousal of high intensity clonic epileptiform states, where the task of separating movement artifact from cortical activity is a work in progress.

For eye artifacts, visual inspection can recognise most types, and we have chosen not to use automated software-removal algorithms to avoid confusion with atypical frontal delta and ISW activity seen in this form of meditation. Visual inspection confirms that the bulk of the frontal activity is quite different to eye-blink artifacts, or lateral eye tracking; for example, if it does occur it is far slower than eye blinks, and is mostly not restricted to frontal EEG sites. However, some cases remain where localised frontal activity may be affected by eye-tracking; these we exclude from detailed ISW source analysis, focusing instead on those cases where ISWs develop sustained high intensities and clear rhythmicity, across multiple electrode sites. In a few cases, some meditators display eye-flutter due to over-concentration, and since spike-wave activity is sometimes also observed, this flutter may be related to eyelid myoclonia (Joshi and Patrick, 2007) seen in absence epilepsy. If it does occur, flutter usually settles down during a recording, but we have found that soft cotton-wool pads held gently in place on the eyelid by lightweight spectacles effectively damps down the physical effects of such activity.

On any remaining occasions where doubt remains about artifacts, those sections are excluded, and all EEG activity analysed in this paper comes from uninterrupted sections to avoid problems of interpolation. The fact that recognisable themes and patterns of cortical sources are found in recordings carried out over several years, using two different amplifiers, with some subjects re-recorded after intervals of 1-3 years, adds to our confidence that we are dealing with patterns of cortical activity inexplicable by artifacts or flawed methodology.

## 3. RESULTS

Table 1 is an overview of EEG recordings of 29 subjects during 2014-18, most carried out during 10-day intensive meditation retreats. Subjects are listed in order of years of meditation experience (subject numbers in the left-hand column represent the order in which subjects were recorded). Two recordings were excluded due to impedance problems leaving 27 subjects, six of whom were re-recorded after intervals of 1-3 years giving 35 independent recordings. Three paroxysmal features were readily recognised in the recordings: spindles, infraslow waves (ISWs) and spike-waves, each accorded a simple measure of feature strengths from visual inspection (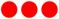 very high, 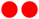 high,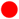 weak/moderate).

**Table 1.**
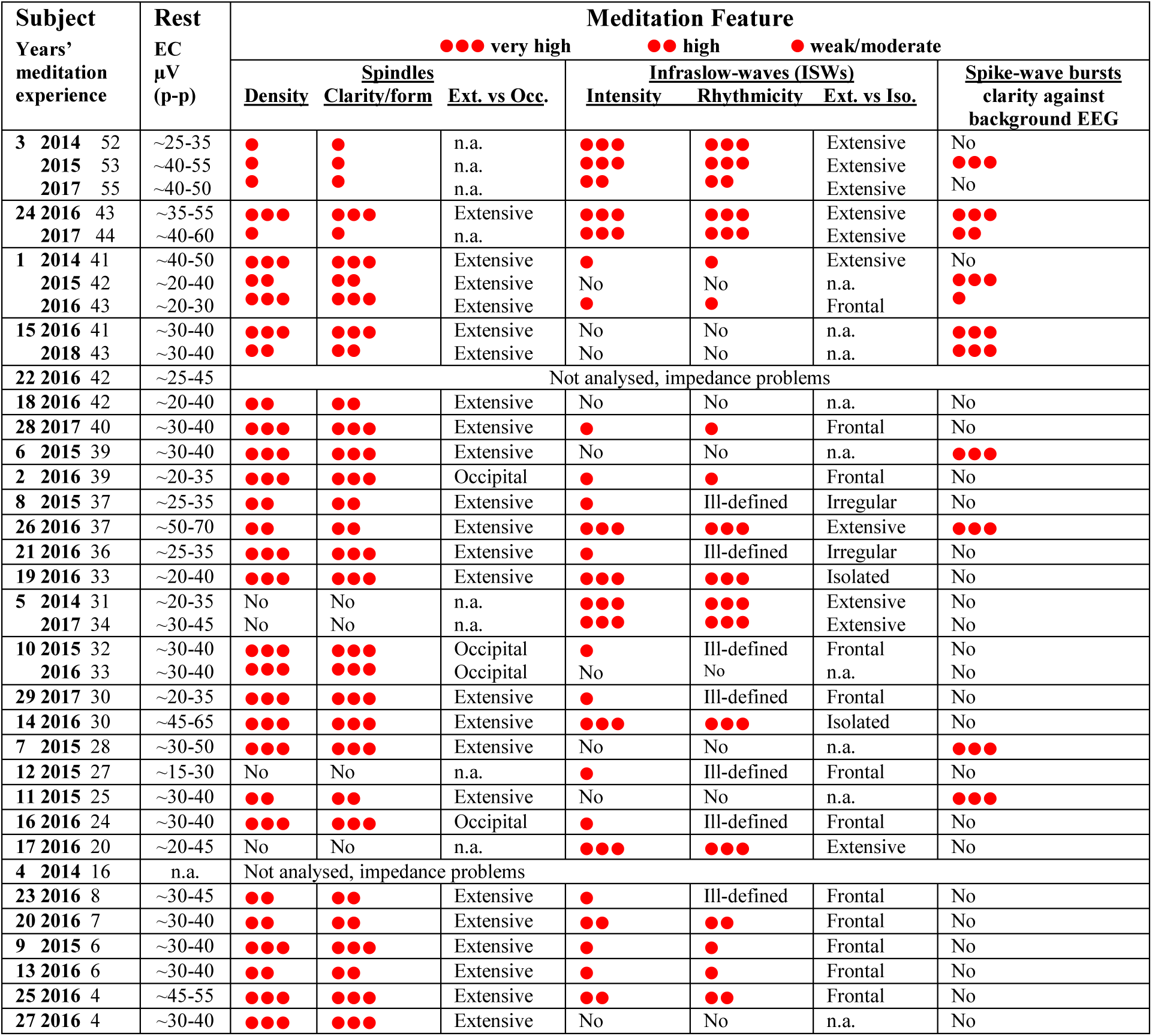
Study **p**articipants: 29 subjects, 35 independent recordings 2014-18. “Extensive” = affecting more than half the electrode sites; p-p = peak to peak. “Very high”, “high”, and “weak/moderate” is a comparative measure of the strength of each feature based on visual inspection.

<u>1. Spindles</u>: 25 subjects (31 recordings) show spindle activity of varying strength. By spindles we refer to “wave-packet” bursts similar to those found in stage-2 nREM sleep, rather than more continuous EEG activity. The 27 recordings marked “high” or “very high” in Table 1 provide the data for a statistical comparison to sleep in both sensor space and source space in Figure 2 below, and the 18 recordings marked “very high” are analysed in source space in Table 2.

**Table 2.** Spindle sources for 18 independent records, 2014-17, with MNI coordinates. SFG, MFG, IFG = superior, middle, inferior frontal gyri; PCL, PCG, PCoG, Rectal = paracentral lobule, postcentral, precognitive, rectal gyri; Pcun, SMG, IPL = precunius, supramarginal gyrus, inferior parietal lobule; pCing, Cing, PHG = postcingulate, cingulate, parahippocampal gyri; ITG, MTG, STG, SMG, Ang, Fus = inferior, middle, superior temporal, supramarginal, angular, fusiform gyri; IOG, MOG, Cun, pCun, Fus, Ling = inferior, middle occipital gyri, cuneus, precuneus, fusiform and lingual gyri.

| Subject<br>Year<br>60 sec<br>samples | B10<br>SFG<br>(Frontal lobe) | B11/B47<br>MFG SFG<br>IFG Rectal<br>(Frontal lobe) | B6/B4/B3/B5<br>SFG PCL<br>PCG PCoG<br>(Frontal lobe) | B7/B40/B19<br>pCun SMG IPL<br>Subgyral<br>(Parietal lobe) | B30/B31<br>pCing Cing<br>PHG<br>(Limbic lobe) | B20/21/22/37/39/40/42<br>ITG MTG STG<br>SMG Ang Fus<br>(Temporal lobe) | B17/B18/B19<br>IOccG MOccG Cun<br>pCun Fus Ling<br>(Occipital lobe) |
| --- | --- | --- | --- | --- | --- | --- | --- |
| 1. 2014 | 19.1% B10 SFG $\pm 5$ 65 -5 | | | | | 8.6% B40 SMG 60 -55 20<br>8.5% B39 MTG -45 -80 20 | 7.3% B18 Ling 5 -90 -20<br>6.5% B19 Fus 40 -65 -20 |
| 1. 2016 |  |  |  | 13.9% B40 SMG 55 -50 30 |  | 10.2% B20 Fus 55 -40 -25 | 11.1% B18 Cun -15 -100 15<br>8.6% B19 Fus 35 -70 -20<br>6.2% B19 MOG 30 -95 10 |
| 2. 2016 |  |  | 16.5% B4 PCG -10 -40 60 | 9.7% B40 Subgyral -35 -45 35<br>7.0% B40 IPL -35 -55 40 | 10.4% B30 pCing 20 -55 5 |  | 6.4% B31 Pcun -5 -70 20 |
| 6. 2015 |  | 10.9% B11 Rectal 5 50 -25<br>8.5% B11 SFG 5 65 -10 | 8.9% B5 PCL 0 -35 60<br>9.4% B3 PCG -20 -40 70 | 12.3% B40 SMG -60 -55 25 |  |  |  |
| 7. 2015 |  |  |  | 15.2% B7 Pcun 10 -50 55<br>11.9% B7 Pcun -5 -70 35<br>10.2% B7 Pcun -10 -65 40<br>(extending to PCG) |  | 6.4% B22 MTG 65 -50 -10<br>6.3% B39 STG 55 -60 30 |  |
| 9. 2015 | | 8.4% B11 MFG $\pm 5$ 65 -15 | | | | | 11.0% B17 Cun $\pm 5$ -100 -5<br>9.7% B18 Ling 10 -100 -10<br>5.9% B17 Ling 15 -95 -5<br>8.0% B18 MOG 15 -90 15<br>7.0% B18 Cun 5 -100 15 |
| 10. 2015 |  |  | 9.0% B6 SFG 5 5 70<br>3.5% B6 SFG -5 -5 70 |  |  | 27.6% B37 Fus 45 -55 -20<br>9.9% B40 SMG 55 -50 20 |  |
| 10. 2016 |  |  | 10.5% B6 SFG 5 15 60 |  | 12.3% B31 pCing 20 -65 15 | 9.0% B21 MTG 65 -55 0 | 18.2% B19 Cun -20 -95 25 |
| 14. 2016 |  |  |  | 10.8% B7 Pcun 15 -55 45 | 17.9% B31 Cing -15 -45 25<br>15.6% B30 PHG -10 -45 -5 |  | 5.7% B19 Fus 25 -75 -20 |
| 15. 2016 |  |  | 8.9% B5 PCL 10 -45 60<br>8.1% B6 PCoG -60 -10 40 | 12.1% B7 Pcun 20 -75 40<br>3.4% B40 IPL 40 -45 40 |  | 4.2% B42 STG -65 -30 15 | 13.3% B18 IOG -40 -90 -10 |
| 16. 2016 |  |  |  |  | 8.4% B30 pCing -10 -55 5 |  | 17.0% B19 Fus -25 -55 -15<br>13.8% B19 Cun 25 -85 30<br>10.8% B19 MOG 40 -70 5 |
| 19. 2016 |  |  |  | 11.9% B40 IPL 65 -35 30<br>10.8% B40 IPL 50 -60 40<br>8.4% B19 Pcun 5 -85 40 |  | 11.0% B20 ITG 65 -10 -25<br>7.9% B39 MTG/Ang -40 -75 35 |  |
| 21. 2016 |  |  |  |  | 10.4% B31 Cing 20 -25 40 | 11.7% B22 STG 50 -35 0<br>9.0% B19 MTG 50 -80 10 | 12.3% B19 Cun -15 -95 25<br>6.6% B18 Ling -10 -90 -20 |
| 24. 2016 |  | 5.5% B11 Rectal 5 15 -25<br>5.4% B11 SFG 15 65 -15 |  | 14.9% B7 Pcun -20 -75 40<br>13.0% B7 Pcun -5 -65 40 |  | 11.2% B39 STG 55 -60 25 |  |
| 25. 2016 |  |  |  | 17.6% B19 Pcun -5 -85 40<br>5.4% B19 Pcun -35 -80 40<br>8.6% B40 IPL 50 -50 25 |  | 9.6% B39 MTG -35 -65 25 | 8.8% B19 MOG 35 -90 15 |
| 27. 2016 |  |  |  | 6.9% B40 IPL -60 -40 25 | 20.6% B31 Cing -10 -45 30 | 8.6% B21 MTG -65 -25 -15<br>13.9% B39 MTG/Ang 50 -75 25 |  |
| 28. 2017 |  | 10.6% B47 IFG -50 30 -10<br>5.7% B11 MFG 30 35 -20 |  |  |  | 17.1% B20 ITG -65 20 -20<br>16.6% B20 ITG/Uncus -30 -5 -40 |  |
| 29. 2017 | | 5.0% B47 IFG $\pm 45$ 35 -5 | | 7.3% B40 IPL -65 -35 35 | | 6.4% B21 MTG 70 -25 -10<br>7.3% B21 MTG 60 5 -15<br>9.3% B39 STG 55 -60 30 | 14.7% B18 Cun 15 -85 25 |
| ROIs | B10<br>SFG<br>Frontal Lobe<br>z-coord -5 | B11 B47<br>MFG SFG<br>IFG Rectal<br>Frontal Lobe<br>z-coord -10—25 | B6 B4 B3 B5<br>SFG PCG PCL<br>PCoG<br>Frontal Lobe<br>z-coord +40—+70 | B7 B40 B19<br>Pcun SMG<br>IPL Subgyral<br>Parietal Lobe<br>z-coord +25—+55 | B30 B31<br>Cing pCing PHG<br>Limbic Lobe | B20/B21/B22/B37/B39/B40<br>/B42<br>ITG MTG STG<br>SMG Ang Fus<br>Temporal Lobe | B17/B18/B19<br>IOccG / MOccG<br>Cun pCun<br>Fus Ling<br>Occipital Lobe |
| Sub-Totals | 19.1% | 60.0% | 74.8% | 211.3% | 95.5% | 230.5% | 208.9% |
| Norm. 100% | 2.1% | 6.7% | 8.3% | 23.5% | 10.6% | 25.6% | 23.2% |
| ROIs Norm. to 100% | B10 B11 B47 Frontal 8.8% |  | B6/B4/B3/B5 Frontal 8.3% | B7/B40/B19 Parietal 23.5% | Limbic 10.6% | Temporal 25.6% | Occipital 23.2% |
|  |  |  | DORSAL PATHWAY |  | VENTRAL PATHWAY |  |  |

<u>2. Infraslow waves (ISWs)</u>: 21 subjects (26 recordings) show varying levels of ISW activity. Those marked “very high” in intensity and rhythmicity and extensive across the head, develop as long trains of well-defined near-sinusoidal ISWs dominating the EEG, and constitute the data for feature characteristics in Figure 7, and analysis in source space in Table 3, below. Such powerful and clearly defined ISW activity is well-suited for cortical source reconstruction. Two other subjects, 14 and 19, also show very strong and well-defined ISWs, but of a quite different nature; isolated, with long silences between, and affecting only frontal and temporal sites; we suspect a different underlying mechanism for these, and reserve their analysis to a future paper.

**Table 3.** ISW Cortical sources for 7 recordings, with epoch lengths 16 or 32 secs according to segment length. MFG, SFG, PCG, MTG, ITG, STG, MOG, IOG, PCoG, PCG = medial frontal, superior frontal, postcentral, middle temporal, inferior temporal, superior temporal, middle occipital, inferior occipital, precognitive and postcentral gyri; PCL = paracentral lobule; pCun = precunius; Cun = cuneus. Each contributing source is listed with its MNI xyz coordinates.

| **Subject**<br>**Individual source contributions normalised to 50%** | **B10**<br>**MFG SFG**<br>**(frontal lobe)**<br>**X +ve/-ve = R/L** | **B11**<br>**MFG SFG**<br>**(frontal lobe)**<br>**X +ve/-ve = R/L** | **B9**<br>**SFG MFG**<br>**(frontal lobe)**<br>**X +ve/-ve = R/L** | **B6**<br>**MFG SFG**<br>**PCL PCoG**<br>**(frontal lobe)**<br>**X +ve/-ve = R/L** | **B5**<br>**PCG**<br>**(parietal lobe)** | **B7**<br>**PCG pCun**<br>**(parietal lobe)** | **B20/B21/B22/**<br>**B37**<br>**MTG ITG STG**<br>**(temporal lobe)**<br>**X +ve/-ve = R/L** | **B18/B19**<br>**MOG / IOG**<br>**Cun**<br>**(occipital lobe)**<br>**X +ve/-ve = R/L** |
| --- | --- | --- | --- | --- | --- | --- | --- | --- |
| **3. 2015**<br>0.032-150 Hz<br>170s segment<br>16s epochs | **9.0%** B10 MFG<br>35 60 -5 | **5.1%** B11 MFG<br>10 65 -15 |  |  |  | **26.9%** B7 PCG<br>-5 -55 70 |  | **9.0%** B18 MOG<br>20 -100 0 |
| **5. 2017**<br>0.032-150 Hz<br>360s segment<br>32s epochs |  |  |  | **27.9%** B6 MFG<br>5 -15 70<br>**12.9%** B6 MFG<br>5 -25 70<br>**9.2%** B6 MFG<br>-5 -20 70 |  |  |  |  |
| **5. 2014**<br>0.032-150 Hz<br>600s segment<br>32s epochs |  |  |  | **15.7%** B6 PCL<br>5 -35 70 | **20.9%** B5 PCG<br>5 -45 70 | **7.1%** B7 PCG<br>5 -55 70<br>**4.2%** B7 PCG<br>-5 -55 70 | **2.1%** B37 MTG<br>-60 -50 -10 |  |
| **17. 2016**<br>0.032-70 Hz<br>300s segment<br>32s epochs |  | **10.8%** B11 SFG<br>-25 55 -15 |  |  |  |  | **16.1%** B22 STG<br>-60 -60 15 | **15.1%** B18 Cun<br>±15 -100 15<br>**8.0%** B18 Cun<br>±5 -100 15 |
| **24. 2016**<br>0.032-70 Hz<br>343s segment<br>32s epochs | **5.5%** B10 MFG<br>35 60 -10<br>**4.6%** B10 SFG<br>-35 50 30 | **9.7%** B11 SFG<br>10 65 -10<br>**6.6%** B11 SFG<br>15 65 -15 | **2.0%** B9 MFG<br>10 50 40 | **5.5%** B6 SFG<br>-20 10 70<br>**3.0%** B6 SFG<br>-5 5 70<br>**3.1%** B6 PCoG<br>25 -15 70 |  |  |  | **10.0%** B18 IOG<br>-40 -90 -5 |
| **24. 2017**<br>0.032-150 Hz<br>400s segment<br>32s epochs |  | **0.7%** B11 MFG<br>5 65 -15<br>**5.2%** B11 MFG<br>-40 55 -10 |  | **6.5%** B6 MFG<br>5 -25 70<br>**4.5%** B6 MFG<br>-5 -30 70 |  | **8.0%** B7 PCG<br>5 -55 70 | **7.0%** B21 MTG<br>65 -55 0 | **18.1%** B19 MOG<br>-40 -90 -5<br>Extends through temp G<br>to ~ -65 -10 -15 |
| **26. 2016**<br>0.032-70 Hz<br>203s segment<br>16s epochs | **1.5%** B10 MFG<br>-30 55 -10 | **12.0%** B11 SFG<br>-20 65 -10 | **1.8%** B9 SFG<br>10 50 35<br>**9.7%** B9 SFG<br>±10 55 40 |  |  |  | **15.7%** B20 ITG<br>-65 -25 -20 | **9.3%** B18 Cun<br>±5 -100 15 |
| **TOTALs** | **20.6%** | **50.1%** | **13.5%** | **88.3%** | **20.9%** | **46.2%** | **40.9%** | **69.5%** |
| Normalised to 100% | **5.9%** | **14.3%** | **3.9%** | **25.2%** | **6.0%** | **13.2%** | **11.7%** | **19.8%** |
| **ROIs**<br>Normalised to 100% | **B10/B11/B9 Frontal**<br>**24.1%** |  |  | **B6 Frontal**<br>**midline vertex**<br>**25.2%** | **B5/B7**<br>**Parietal midline vertex**<br>**19.2%** |  | **B20/B21/B22/B37**<br>**Temporal**<br>**11.7%** | **B18/B19**<br>**Occipital**<br>**19.8%** |

<u>3. Spike-wave bursts</u>: 8 subjects (11 independent records) show clearly defined and occipital spike-wave bursts.

The presence of persistent and strong ISWs and spindles invites comparison to nREM sleep, anaesthesia and coma, while the occurrence of spike-waves is reminiscent of absence epilepsy. In this section each theme is taken in turn with the findings considered as a whole in the final Discussion.

### 3.1. SPINDLES

Figure 1 shows three examples of meditation spindles using a 5.3-15 Hz bandpass to reduce background ISWs and higher frequency beta and gamma activity. Superficially, spindles appear as symmetrical packet-like bursts of rhythmic activity in the EEG, similar to those in sleep, but more prolific which makes their recognition easier than in polysomnography. Some subjects show extensive and widespread spindles (e.g. middle panel), others less extensive, but all recordings show involvement of occipital sites. The upper panel and enlarged extract shows very well-defined occipital spindles, with a symmetrical waxing/waning morphology. The middle panel shows widespread spindling across most electrode sites, and illustrates how in some cases spindles develop into spike-waves (Leresche *et al.*, 2012), suppressing spindle activity, in this case at occipital sites with a frequency of 5.37 Hz for the 3-sec yellow-highlighted segment. The lower panel again shows widespread spindling, although not so extensive.

**Figure 1.**
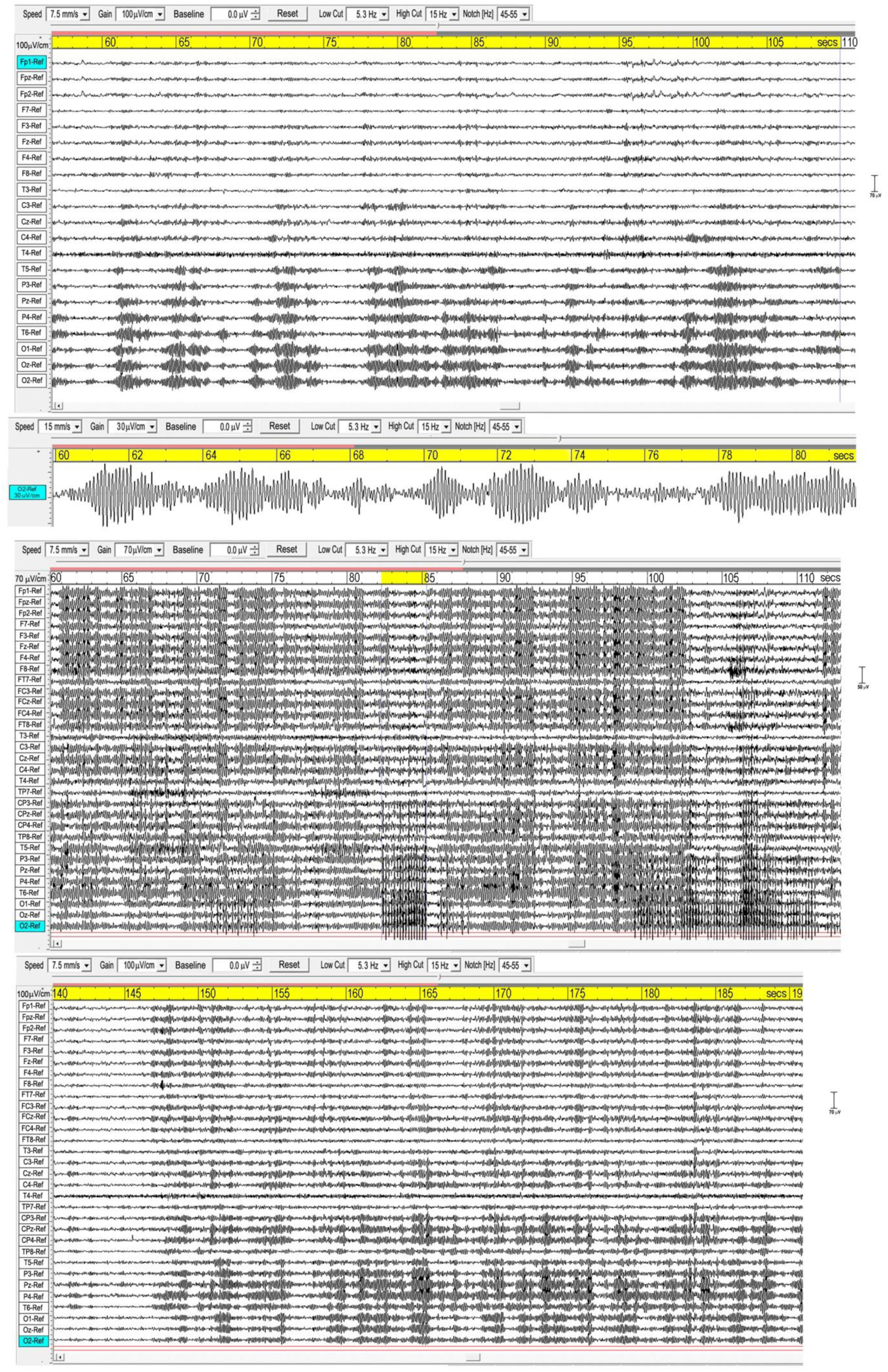
Three examples of spindling. The upper panel is from a recording of subject 16, 2016; the middle panel subject 1, 2015; and the lower panel subject 7, 2015. Bandpass 5.3-15 Hz.

**Figure 2.**
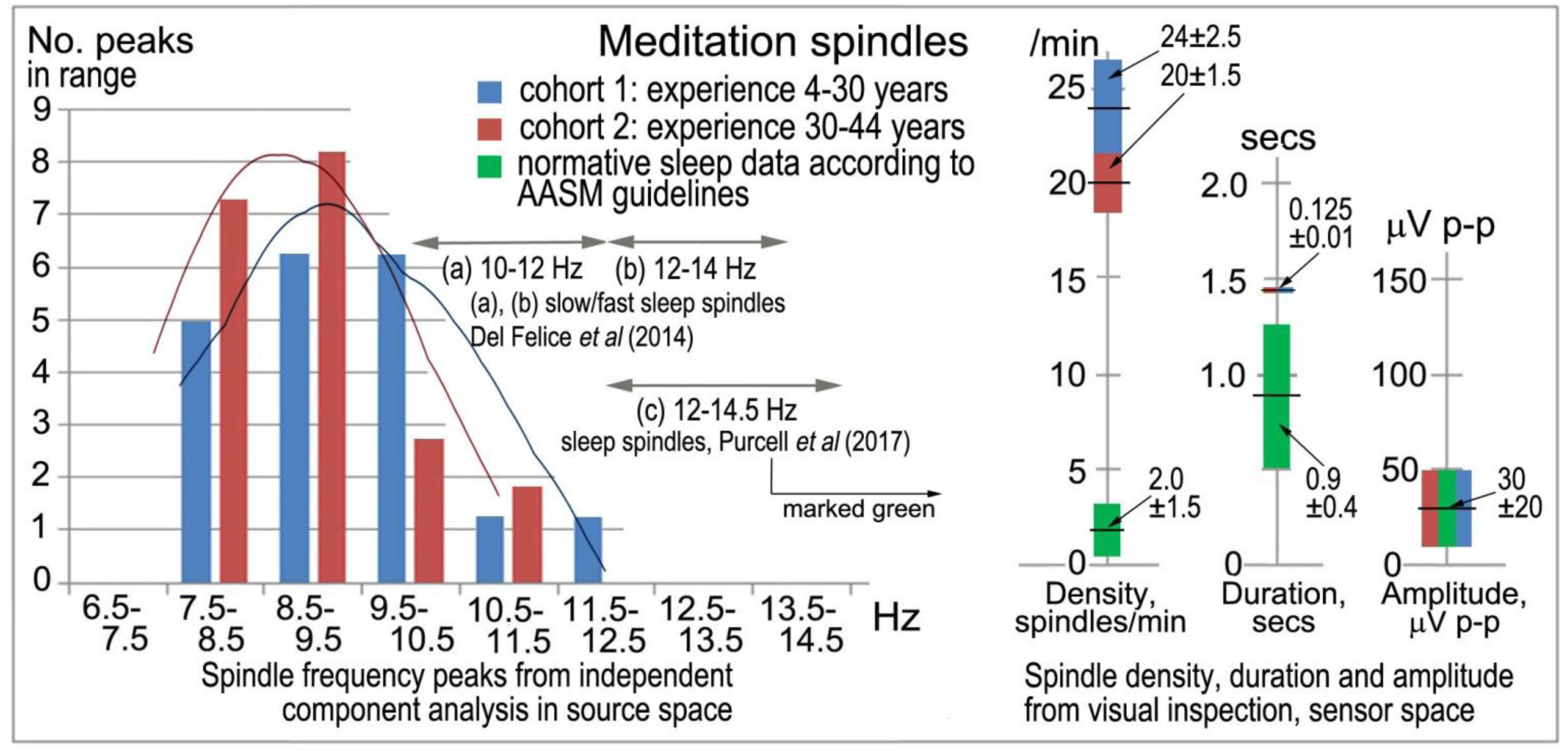
Meditation spindles compared to those in stage-2 nREM sleep for 27 independent recordings. The bar charts (left) show spindle frequencies from an independent component (IC) spectral analysis in source space of 60-sec segments from each recording. The data are split into two cohorts of meditation experience, 4-30 years and 30-44 years; the former yielding 16 spectral peaks and the latter 22 peaks, normalised to 20 for each cohort in the plot. At right are spindle density, duration and amplitude in sensor space from visual inspection of <u>N</u>=30 consecutive spindles in each record (spindles at least 3x the inter-spindle background amplitude; total <u>N</u>=810 spindles), compared to values typical of sleep. American Academy of Sleep Medicine guidelines (AASM, 2017) were followed for visual inspection.

Sleep spindles in the sigma band (9-16 Hz) are well-researched in polysomnography (e.g. Purcell *et al*’s., 2017, metastudy), and Figure 2 compares spindle parameters between the two modalities.

Although the sample size is not large, the spindle frequency bar charts in source space for the two cohorts of meditation experience appear different. The curves of best fit in Figure 2 are non-parametric plots of the distribution of spindle frequencies for the two groups, obtained using a kernel density estimator (implemented using Stata’s kdensity command, with an imposed bandwidth of 0.8). A linear probability model in which a dummy is regressed for having a spindle frequency <10 Hz against a dummy identifying each of the two cohorts, shows that meditators with >30 years’ experience are statistically more likely to experience spindle frequencies below 10 Hz than meditators with <30 years’ experience. Meditators with more than 30 years’ experience are 38% more likely to experience spindle frequencies below 10 Hz (p=0.092).

Spindle density, duration and amplitude in sensor space compared to sleep are shown in Figure 2 (right). While there is no difference in amplitude, meditation spindle density is an order of magnitude greater than in sleep, which greatly aids the credibility of analysis; the difference between the two meditation cohorts is also statistically significant (p<0.1) suggesting spindle density lowers slightly with longer experience. Spindle duration is also significantly greater (∼7x) in meditation than in sleep (p<0.1).

Underlying cortical sources were computed for the 18 recordings showing spindles with the highest density and clarity of form (Table 1), using eLoreta to identify the strongest ICs accounting for at least 50% of the signals’ variance of each 60 sec sample, with total variance normalised to 50%. Figure 3 is an example for subject 14, 2016, and Table 2 summarises the results for all 18 independent records, with columns 2-8 listing cortical sources with MNI (Montreal Neurological Institute) xyz coordinates.

**Figure 3.**
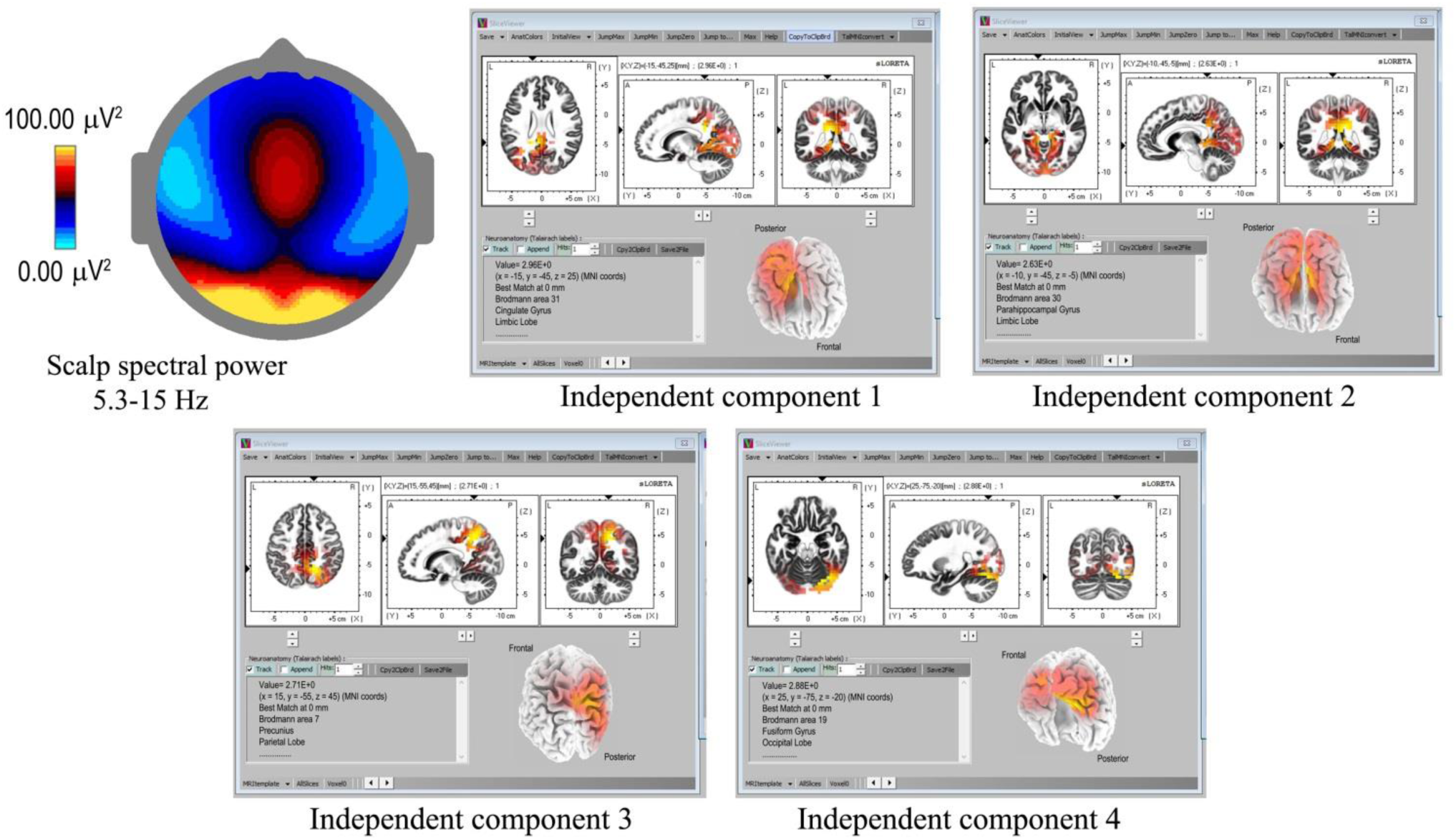
Spindle sources computed using eLoreta for a 60-sec sample of spindles (subject 14, 2016), bandwidth 5.3-15 Hz, 4-second epochs. The scalp mean spectral power distribution at upper-left shows maximum intensity in occipital regions, with some extension along the midline. The two strongest sources (ICs 1, 2) are limbic, at Brodmann sites B31 and B30, the cingulate and parahippocampal gyri respectively; with IC3 and IC4 at B7 (temporal) precunius, and B19 (occipital) the fusiform gyrus.

The bottom portion of Table 2 summarises the spindle cortical sources into dominant regions of interest (ROIs). The anterior-most frontal sources are in Brodmann areas B10, B11 and B47, amounting to 8.8% of total signals’ variance; frontal sources closer to the vertex at Brodmann areas B6, B4, B3, B5 amount to 8.3%; parietal sources at Brodmann B7, B40, B19, 23.5%; limbic sources at Brodmann B30, B31, 10.6%; temporal sources, 25.6%; and occipital sources, 23.2%. The significance of the lower labels dorsal and ventral streams will be considered in the Discussion.

### 3.2. INFRASLOW-WAVE (ISW) ACTIVITY

In sleep studies, slow waves (SWs) are considered to reflect bistability of cortical neurons undergoing slow oscillation (<1 Hz) between two distinct states, and are discriminated from paroxysmal discharges by their symmetrical and rhythmic shape. Meditation slow waves are also rhythmic, quite different to paroxysmal discharges, with frequencies well below 1 Hz distinguishing them from 1.0-4.0 Hz delta activity, in fact sufficiently lower to justify the label infraslow waves (ISW). Figures 4 and 5 show six independent recordings illustrating main features. EEG electrode sites are labelled left, from frontal (F) sites at the top, to occipital (O) sites at the bottom (T, C, P denote temporal, central and parietal areas). The top bar shows time in secs. Figure 4, top panel (subject 5, 2014), shows intense ISWs at frontal, occipital and central-temporal sites. The inset scalp intensity distributions correspond to the start and end points of the yellow-highlighted interval in the time bar, showing a pattern of alternating ISW inhibition-excitation, in this example reaching 1350 µV p-p at CPz.

**Figure 4.**
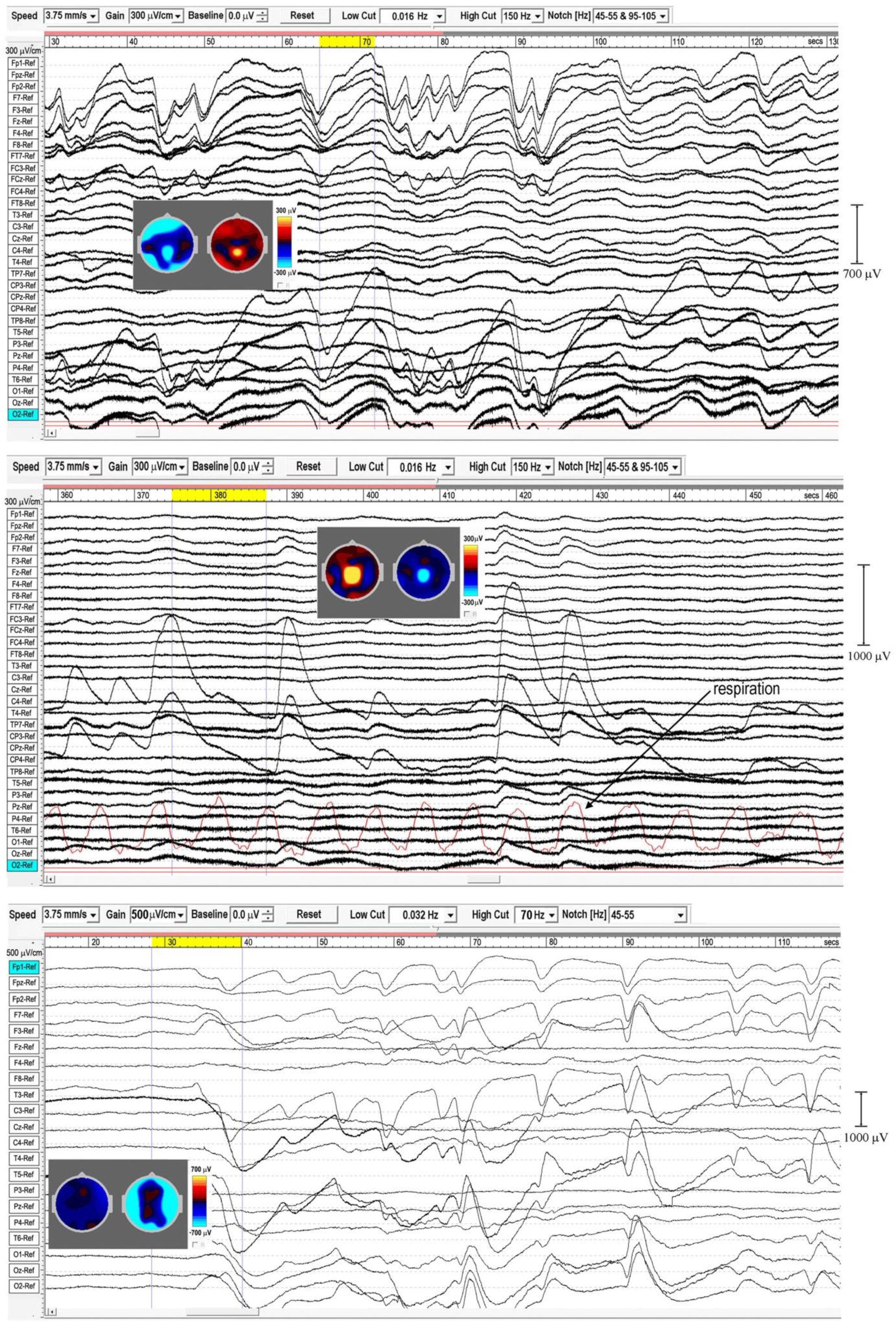
Extensive and powerful infraslow waves during Samatha meditation. Top panel subject 5, 2014; middle panel subject 5, 2017; bottom panel subject 17, 2016. The inset scalp intensity maps correspond to the start and end points of the yellow-highlighted intervals.

**Figure 5.**
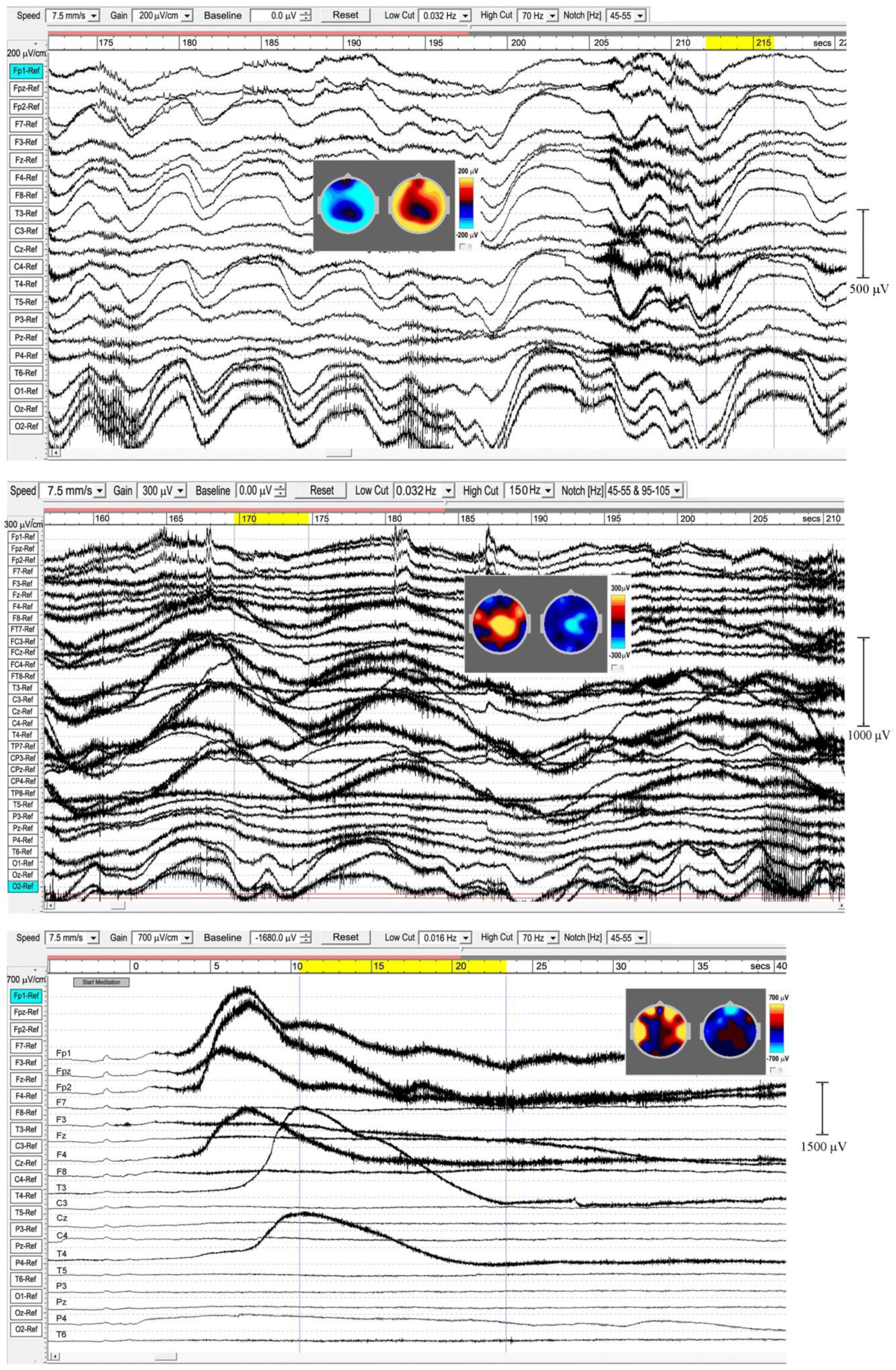
Extensive and powerful infraslow waves. Top panel subject 24, 2016; middle panel subject 24, 2017; bottom panel subject 19, 2016. Note the posterior spike-wave bursts for subject 24, and the isolated extremely high voltage ISW shown by subject 19.

The middle panel is the same subject re-recorded in 2017, with more experience of jhāna meditation, showing highly focused activity near the vertex at Cz and CPz (highlighted by the inset scalp maps) with p-p values >1500 µV, and much reduced frontal and occipital activity. The form of the ISWs is also different; a more rapid +ve onset, and slower recovery reminiscent of patterns for relaxation oscillators.

The rate of rise for the ISW at 418 secs (Figure 4) reaches ∼1400 µV/sec; the red trace is respiration, measured by an induction-loop chest belt. The bottom panel, subject 17, 2016, illustrates how ISW onset in some cases can be very rapid; in this case widespread ISW activity starts at ∼35 secs into meditation, with an initial massive inhibition at 40 secs, with the inset maps showing a complete annulus of inhibition around central areas. The ensuing ISWs reach remarkable intensities > 2000 µV p-p at times.

The top panel in Figure 5 (subject 24, 2016) shows ISWs particularly strong at occipital sites, with inset maps for the ISW at 212-216 secs again showing an annulus of inhibition-excitation enclosing central areas. This subject also shows brief periods of enhanced gamma activity (e.g. at 206-12 secs), as well as spike-wave bursts lasting 2-6 secs at occipital sites. The middle panel shows the same subject in 2017, now with much stronger ISW activity at sites around the vertex (see inset scalp maps) compared to 2016. Occipital ISWs are still present, also significant gamma activity, and spike-wave bursts at occipital sites as in 2016. In contrast, the lower panel (subject 19, 2016) is one of two examples (noted earlier, Table 1) which we believe illustrate a different mechanism, to be explored separately, of isolated ISWs with longer periods of recovery and relative “silences” between. This subject shows fast responsiveness and ISW onset similar to subject 17 in Figure 4, with a massive positive ISW appearing just 5 secs after starting meditation, reaching 2400 µV at Fp2, one of the highest levels we have recorded; travelling ∼3.6 secs later to temporal sites T3 and T4. For this subject, the intense ISWs are accompanied by strong increases in the gamma band. The scalp maps for the segment 10.5-23.3 secs show the familiar alternation of excitation-inhibition, which in this case is focused at frontal and temporal sites.

When respiration has been measured, visual inspection suggests a close relationship between ISW and respiration frequencies; e.g., subject 5, 2017, middle panel Figure 4, showed a mean ISW period of 9.59 ± 0.66 secs, and mean respiration period for the same segment of 9.88 ± 0.33 secs. However, a more detailed correlation analysis has not yet been possible due to software limitations in WinEEG.

#### 3.2.1. ISW Statistics

The left-hand bars in Figure 6 summarise statistics in sensor space from >600 ISWs from 8 independent recordings of 5 subjects, 2014-17, rated “very high” in intensity and rhythmicity in Table 1. Sections showing at least 3 successive ISWs were examined in turn, with periods between successive +ve to +ve, or –ve to –ve peaks measured, according to whether +ve or –ve peaks were dominant (Figure 4, middle panel, e.g., shows +ve-dominant ISW peaks). Peak-to-peak voltages from either a +ve peak to the following minimum, or from a –ve peak to the following maximum were measured giving a mean p-p ISW amplitude of 417±69 µV, with individual SWs ranging from ≲100 to 2255 µV p-p.

**Figure 6.**
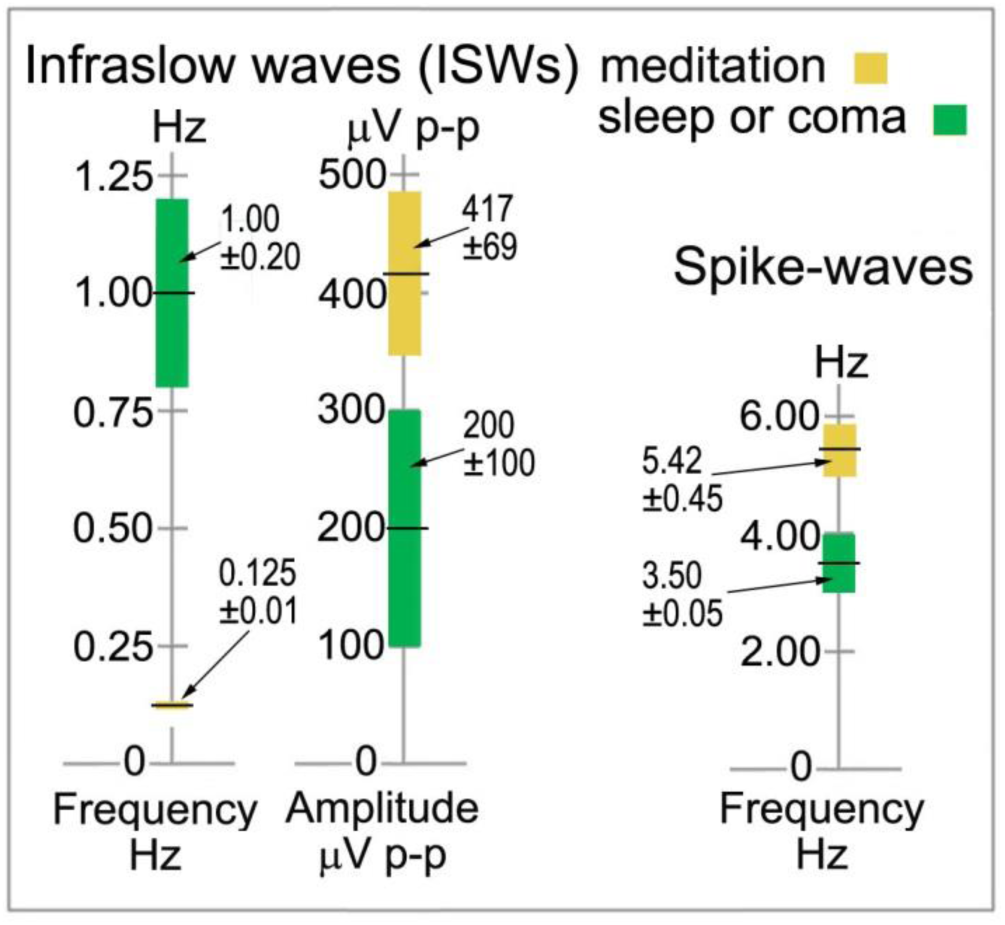
Infraslow wave and spike-wave statistics in sensor space compared to sleep and absence epilepsy

**Figure 7.**
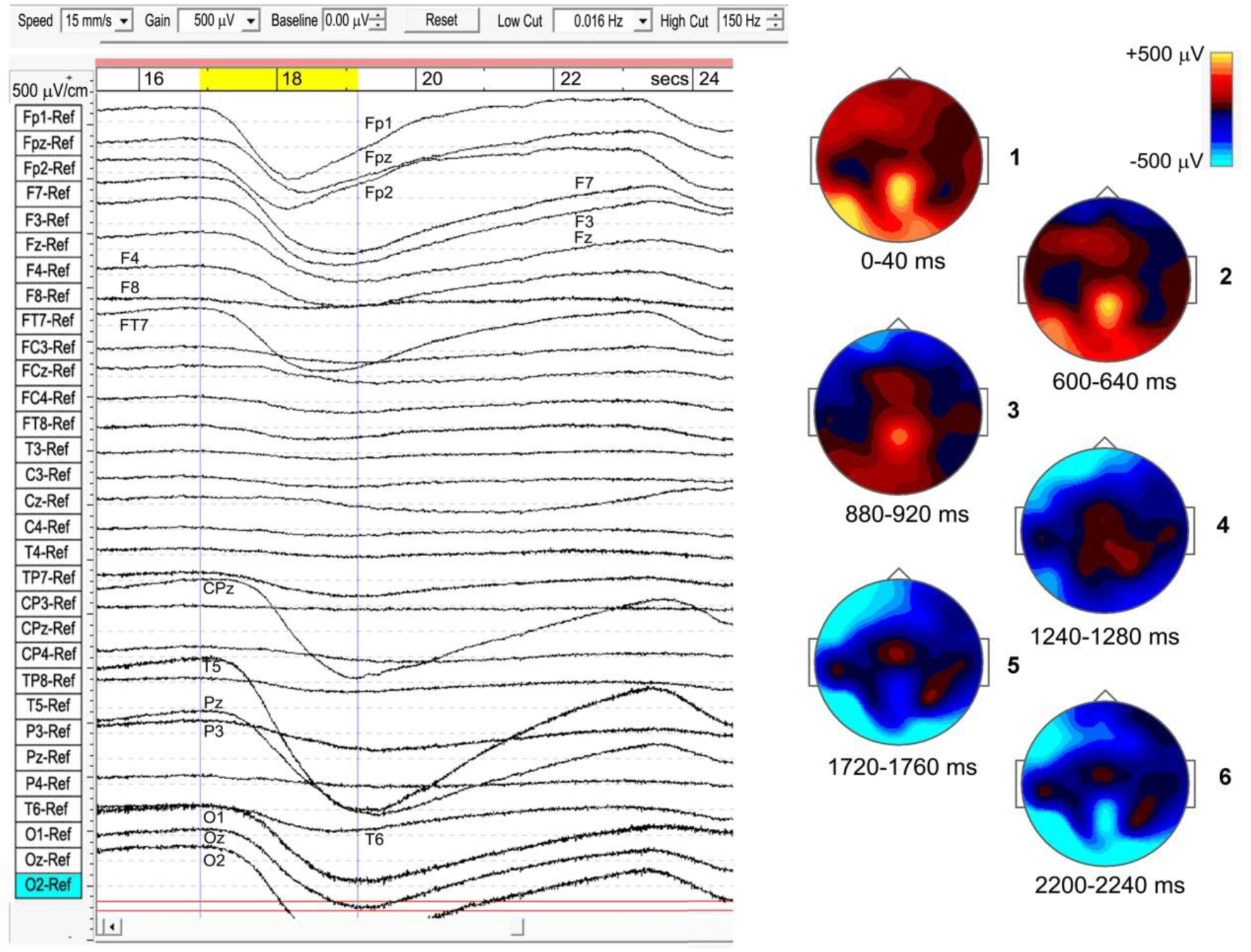
Example of a travelling infraslow wave (subject 5, 2014), bandpass 0.016-150 Hz.

The 8 recordings yielded a mean ISW period of 8.36±0.60 secs, corresponding to a mean frequency of 0.125±0.010 Hz. The corresponding values for sleep SWs are 1.00±0.20 Hz and 200±100 µV (Libenson, 2012; Sutter and Kaplan, 2012) with the differences statistically significant at p<0.1.

#### 3.2.2 Travelling ISWs

The strongest examples of ISWs noted in Table 1 all show a “travelling” nature to varying degrees, as do some sleep slow waves. In some cases the meditational ISWs appear nearly simultaneous for 10’s of seconds across wide areas, while other sections of the same recording might show frontal sites leading or lagging posterior or temporal sites. As far as we can determine there is no obvious relation to the stage of a subject’s meditation. Subject 5, 2014 (top panel, Figure 4), sustained strong and well-defined travelling ISWs, 300-700 µV p-p, for over 20 mins, and provides the clearest example, shown in Figure 7 where the scalp intensity maps 1-6 trace the development of the -ve ISW peak across a 2.25-second interval (highlighted yellow, top bar). Note the localised midline peak, posterior to the vertex, extending occipitally. Also, the considerable, and typical, broadening of the ISW as it reaches occipital areas. Scalp map 1 shows the initial +ve phase of the wave, strongest at left-temporal site T5, and midline central-parietal CPz, less intense frontally. The –ve phase then develops frontally, extending left-frontally, before travelling to occipital areas (mainly left), as in maps 2-6.

Maps 1 and 6 in Figure 7 correspond roughly to the excitation-inhibition peaks. The localised midline peak, just posterior to the vertex, is also observed for other subjects who show strongly developed and extensive ISWs. Although this is surface EEG activity, the near-vertex peak will be seen to recur in source space.

Although for most meditators the sites of activity remain fairly constant during a recording (in this example, EEG sites Fp1, Fpz, Fp2, F7, F3, Fz, F4, FT7, CPz, T5, Pz, O1, Oz and O2), transit times vary considerably. In this case, the ISW in Figure 6 shows a transit time of the –ve peak, front to rear, ∼1200 ms. For 20 successive ISWs from this same subject, transit times varied widely, with means over 20 mins of: frontal (Fp1) to occipital (O1/O2), 593±300 ms; frontal (Fp1) to temporal (T5), 723±230 ms; and frontal (Fp1) to central-midline (CPz), 1278±340 ms. The mean front to rear transit time ∼600 ms corresponds to a mean transit speed ∼50 cm/s across the head, significantly slower than typical values ∼1.2-7.0 m/s for sleep SWs (Massimini *et al*., 2004).

#### 3.2.3 Rhythmic Excitation-Inhibition

Meditation ISWs are not random, exhibiting rhythmic patterns of powerful excitation-inhibition, as in Figures 4 and 5. In some cases this activity forms an almost complete annulus of excitation-inhibition around less affected central areas (e.g. subject 17, 2016, Figure 4; and subject 24, 2016, Figure 5). For other subjects the annulus is only partial, as for subject 5, 2014, Figure 4; and subject 19, 2016, Figure 5. For subjects 5 and 24, re-recorded in 2017 (Figures 4 and 5 respectively) and both with more experience of jhāna, the annulus is replaced by an intense focus near the vertex.

#### 3.2.4 Cortical Sources

Table 3 summarises a source analysis for 7 independent recordings from the same 5 subjects who show the strongest examples of extensive ISW activity. Subject 5 was recorded twice, in 2014 and 2017, both with the 31-electrode system; and subject 24 also twice, in 2016 and 2017, with the 21- and 31-electrode systems respectively. To study low frequency structure <1 Hz, epoch length is a key variable (see section 2.2), and depending on the segment lengths, epochs of 16 or 32 secs were used. We are therefore confident of capturing frequencies down to 0.03 Hz for five of these recordings (32 sec epoch) and to 0.06 Hz for the other two, particularly given the mean ISW frequency of 0.125±0.010 Hz noted above.

As with spindle analysis, cortical sources were computed using eLoreta for the strongest ICs accounting for at least 50% of the signals’ variance for each sample, then normalised to 50%, and summarised in Table 3. Averaged across all 7 recordings (>2500 secs of strong and persistent ISWs), sources are found as follows: frontal sites, Brodmann B10, B11, B9 (24.1% of the total variance across all recordings); frontal midline site B6 just anterior to the vertex (25.2%); parietal midline B5, B7 just posterior to the vertex (19.2%); temporal B20, B21, B22, B37 (11.7%); and occipital sites B18, B19 (19.8%).

These ROIs are summarised visually in the upper part of Figure 8, with 3D source maps to aid visualisation based on superposition of individual components from eLoreta that contribute to each ROI. The dominant ROI is in the midline vertex area, bridging the frontal-parietal junction, accounting for 44.4% of total signals’ variance. The supporting ROIs are frontal sources (24.1%); temporal sources (11.7%), predominantly left and diffuse, merging with, again predominantly left, inferior and middle-occipital sources (10.6%); and finally at far-right the midline cuneus (9.2%).

**Figure 8.**
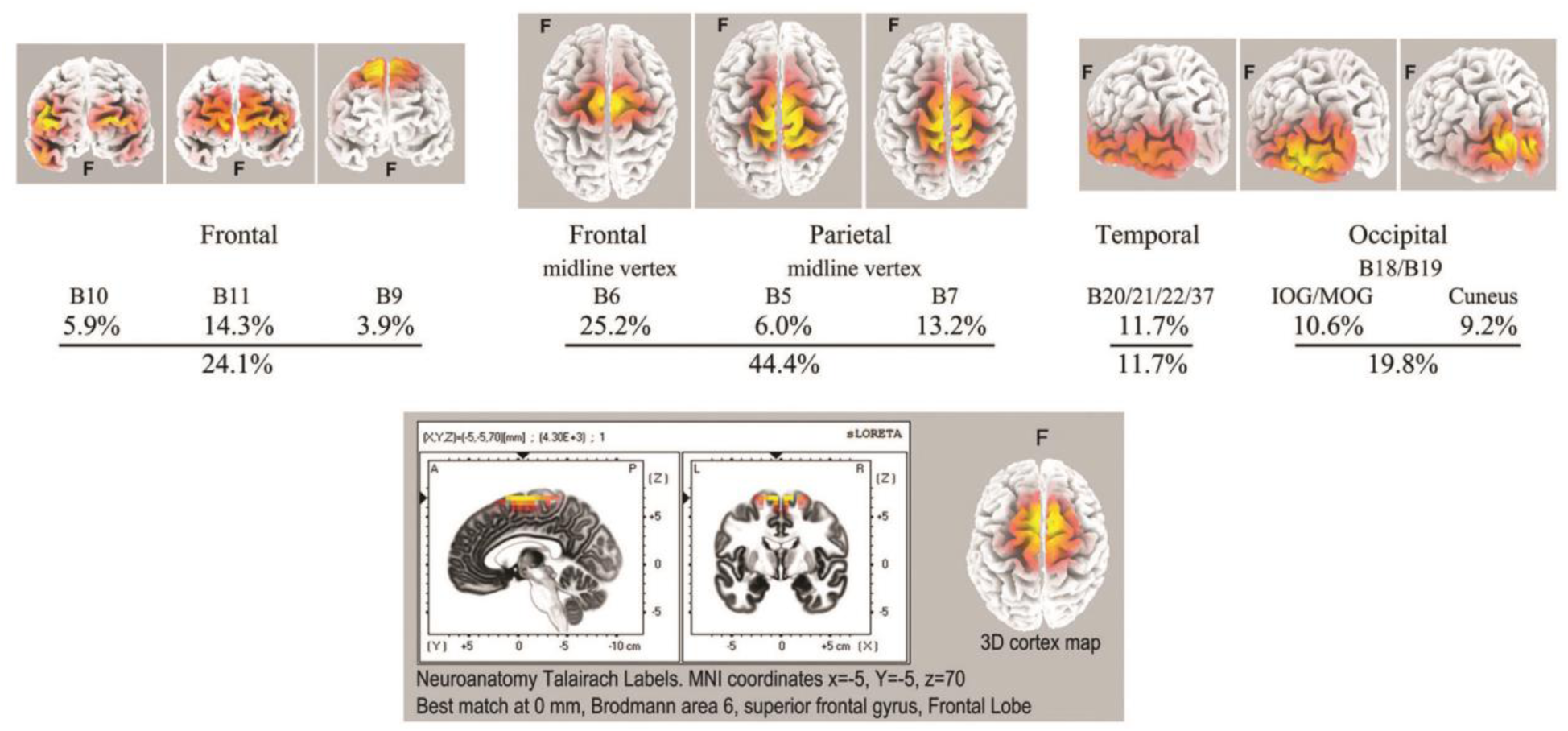
Top: summary of ISW regions of interest from Table 3 (7 independent recordings), with 3D cortical source plots. The fully developed vertex source is illustrated below for subject 5, 2017.

The lower part of Figure 8 illustrates the highly focused vertex activity that develops in deeper jhāna meditation, this example from subject 5, 2017, which we will return to in the Discussion.

#### 3.2.5 An Underlying Slower ISW Component

We have two examples of particularly strong ISW activity that suggest an underlying rhythm even slower than the mean 0.125 Hz ISW frequency noted above (Figure 6). Such very low frequency EEG activity has been little studied in neuroscience, largely due to difficulties with DC drift, confusion with noise and artifacts, as well as the requirement of long records to allow epochs ideally >60 secs. WinEEG imposes a software limitation to spectral analysis below ∼0.02 Hz, so at this point we rely on visual inspection, with spectral analysis as a tentative comparison. Figure 9 shows extracts from subjects 17 (2016) and 5 (2017) with, right, superposed ISWs from longer segments (300 and 600 secs respectively) showing a morphology of rapid leading edge and slower over-shooting recovery. The superpositions show half-periods ∼21 and ∼26 secs respectively, suggesting an underlying very slow ISW frequency ∼0.02 Hz. Bearing in mind software limitations, a spectral analysis of a 600-sec segment from subject 5, using 64-sec epochs and 0.016 Hz low cut, shows some evidence of peaks at 0.02-0.03, and 0.05 Hz.

**Figure 9.**
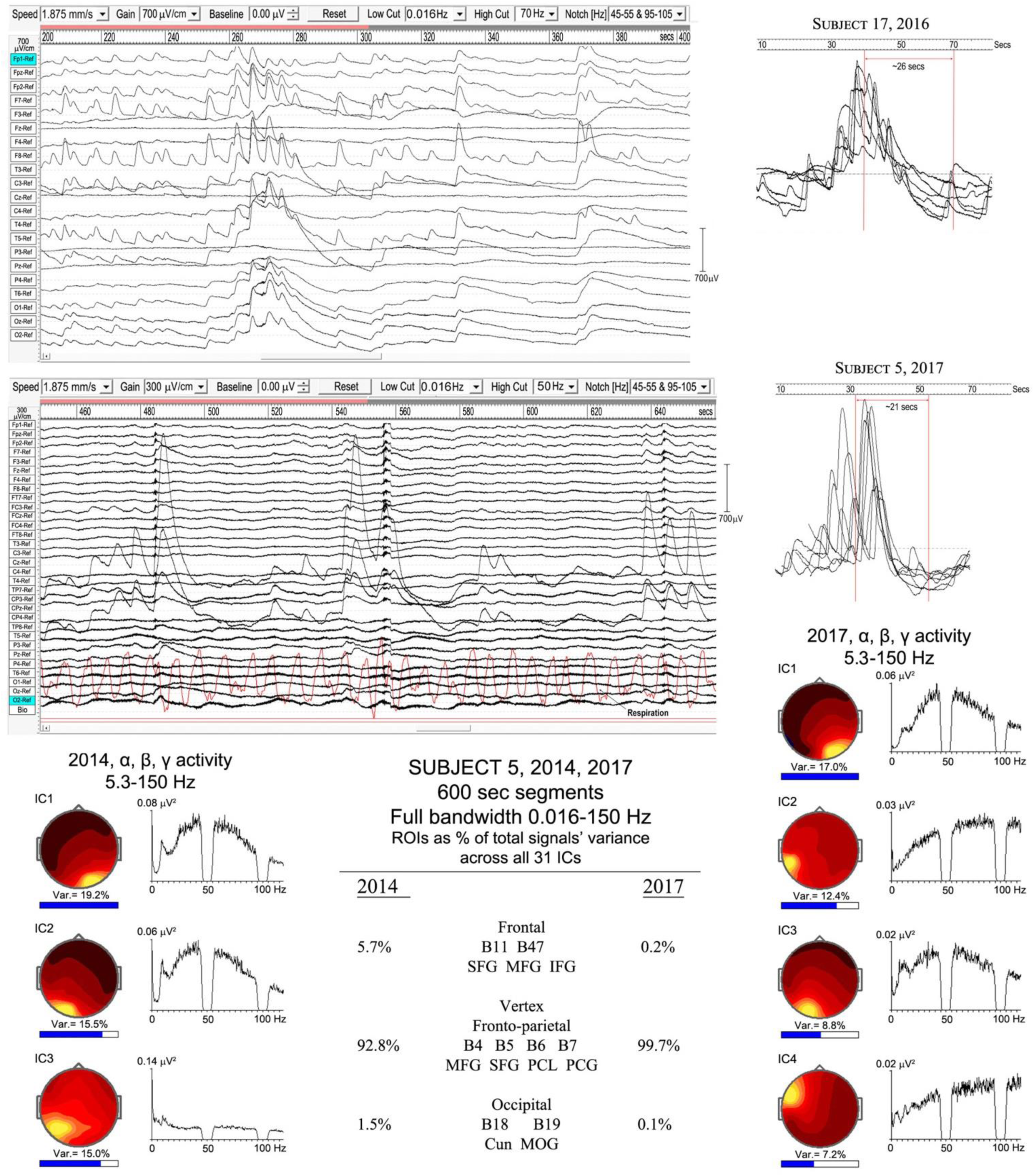
Very slow infraslow-wave (ISW) activity: top panel, subject 17 (2016); middle panel subject 5 (2017). Below is a comparison of the highly focused vertex activity for subject 5 in 2014 and 2017, with the strongest ICs from 600-sec samples shown for each year, with overall percentage contributions of ROIs from all 31 ICs for each year. The IC spectra alongside show higher frequency activity (bandwidth 5.3-150 Hz), revealing broadband gamma with only small residual traces of alpha activity.

The lower part of Figure 9 illustrates the overwhelming dominance of ISW activity near the vertex for subject 5 in 2014 and 2107. The overall percentage contributions of ROIs to the total signals variance calculated from all 31 ICs for each year from 600-sec samples are shown; being 5.7% frontal, 92.8% vertex and 1.1% occipital in 2014; and 0.2% frontal, 99.7% vertex and 0.1% occipital in 2017. To examine what remains apart from ISW activity, the strongest IC spectra were computed using a 5.3-150 Hz bandwidth and are shown at each side, with scalp intensity distributions, revealing broadband gamma activity with only small residual traces of alpha activity, particularly weak in 2017.

### 3.3. SPIKE WAVE AND SEIZURE-LIKE OR EPILEPTIFORM ACTIVITY

#### 3.3.1 Spike-Wave Activity

Spike-waves are regarded as the signature of absence epilepsy (Sadleir *et al*., 2009), and refer to bursts of sharp spikes in the EEG (duration 20-70 ms), each followed by a “recovery” ISW, the spikes repeating at ∼3-4 Hz. The meditational spike waves we observed were quite unexpected, appearing to arise spontaneously with meditators fully alert and unaware of any specific change in subjective experience. Table 1 lists 8 subjects (11 independent recordings) showing brief bursts of occipital spike waves of 3-12 secs duration, similar to absence epilepsy, apart from subject 26 who showed this activity for ∼50 secs. Some meditators also develop spike waves during the deliberate arousal of strong energisation, or *pīti*, the third jhāna factor mentioned in the Introduction. The upper panel of Figure 10 shows 4 examples, with bandpass 0.53-70/150 Hz to minimize background ISWs while retaining high-frequency content. Harmonic structure is apparent in the spectra of the main ICs for the two subjects in the lower part of Figure 10, which also shows spectral intensity distributions. For subject 1, spike wave activity is focused at Brodmann B18, middle occipital gyrus (MOG) left, and cuneus, right; and for subject 7 at Brodmann B17 lingual gyrus left, and Brodmann B19 cuneus right.

**Figure 10.**
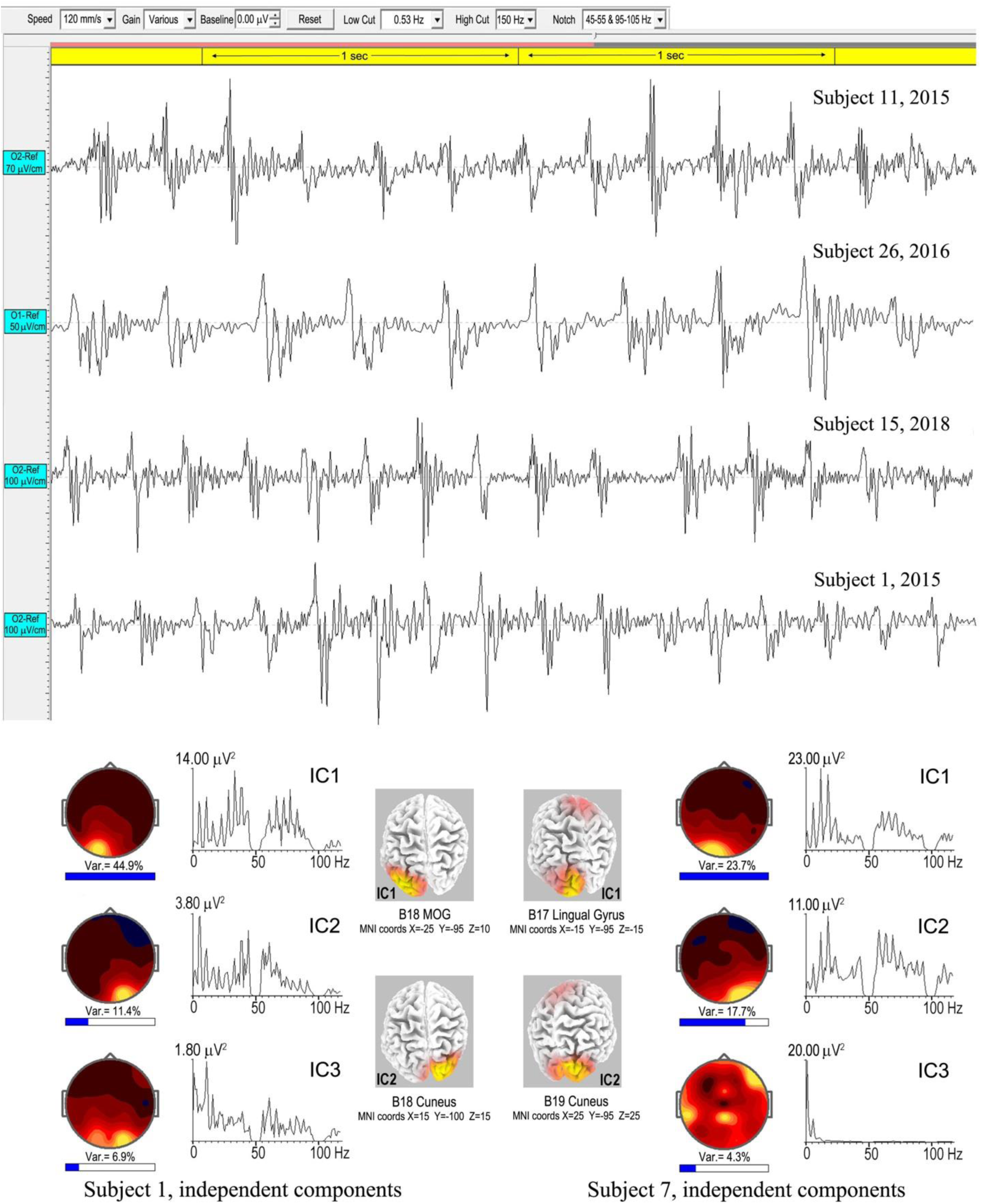
Above are four **e**xamples of spike-wave bursts at occipital sites, using a bandwidth 0.53-70/150 Hz. Top to bottom are excerpts from an 8.6-sec burst, subject 11, 2015; a 50-sec burst, subject 26, 2016; an 8.3-sec burst, subject 15, 2018; and a 3.025 sec burst, subject 1, 2015. Below are the strongest ICs for subjects 1 and 7, from 3.025-sec and 7.0-sec bursts respectively, computed using eLoreta, showing harmonic spectral structure, spectral intensity distributions, and 3D source maps.

Table 4 summarises an eLoreta analysis in source space of spike wave bursts at least 3 secs in duration and well-defined against background activity, from 9 independent recordings. We find harmonic structure in all cases, with a range of spectral peaks for the strongest ICs shown in column 7, with harmonics underlined. Cortical sources with MNI xyz coordinates are in columns 3-6, with mean ROIs shown in the bottom row. Occipital sources overwhelmingly dominate at Brodmann areas B17, B18, B19 (electrode sites O1, Oz and O2 in sensor space), accounting for 88.7% of total signals’ variance, with 9.6% and 1.7% frontal, and frontal-near-vertex, contributions respectively. Since some SW breakthrough remains in some cases, this occipital dominance is likely an underestimate.

**Table 4.** Spike-wave sources from eLoreta for 9 independent records, 2015-18, bandpass 0.53-70/150 Hz, 1-sec epochs. MFG, SFG, IFG = medial, superior and inferior frontal gyri; IOccG, MOccG = inferior and middle occipital gyri; pCun, Cun, Ling = precuneus, cuneus and lingual gyrus. For each subject, the 3-6 strongest ICs were used that accounted for at least 50% of the signals’ variance, then normalised to exactly 50% variance, for each subject. The final totals across all the recordings are normalised to 100%. An IC algorithm was used for subjects 3 and 11 to remove eye-blinks.

| Subject year recorded | Spike-Waves | B10 MFG<br>(frontal lobe) | B11/B47 MFG SFG IFG<br>(frontal lobe) | B6 MFG<br>(frontal lobe) | B17/B18/B19 IOccG MOccG pCun Cun Ling<br>(occipital lobe) | S-W spectral peaks from independent components |
| --- | --- | --- | --- | --- | --- | --- |
| **1. 2015** | 3.025 sec. segment<br>At O1, O2, Oz<br>0.53-150 Hz<br>1-sec epochs |  |  |  | **35.5%** B19 MOccG<br>-25 -95 10<br>**14.5%** B18 Cun<br>15 -100 15 | <u>5.37, 10.74, 16.13, 22.46, 28.32, 33.20, 38.58, 42.97, ... 60.55, 66.41, 71.78, 77.15, 83.01</u> |
| **3. 2015** | 8.2 sec. segment<br>At O1, O2, Oz<br>0.53-150 Hz<br>1-sec epochs |  | **7.1%** B11 MFG<br>-10 65 -15 |  | **23.8%** B17 Ling<br>20 -95 -5<br>**19.1%** B19 MOccG<br>-40 -90 5 | <u>5.36, 10.74, 15.63 Hz</u> |
| **6. 2015** | 12.0 sec. segment<br>At O1, O2, Oz<br>0.53-150 Hz<br>1-sec epochs | **7.9%** B10 MFG<br>10 65 0 |  | **7.9%** B6 MFG<br>5 -30 70 | **17.3%** B19 IOccG<br>-40 -85 -10<br>**9.5%** B18 Cun<br>15 -100 0<br>**7.4%** B19 MOccG<br>45 -85 10 | <u>3.91, 7.81, 11.72, 15.63, 19.53, 23.44, 27.34, 31.25, 32.23, 36.13, 40.04, 43.95, ... 72.27, 76.17, 80.08... Hz</u> |
| **7. 2015** | 7.0 sec. segment<br>At O1, O2, Oz, Cz<br>0.53-150 Hz<br>1-sec epochs |  | **6.4%** B47 IFG<br>-50 45 -10 |  | **25.3%** B17 Ling<br>-15 -95 -15<br>**18.3%** B19 Cun<br>25 -95 25 | <u>5.85, 11.72, 17.58, 23.44, 37.11, 41.99, 58.59, 64.45, 70.31, 76.17, 83.01, 85.94, 88.87, 92.77 ... 105.47, 110.35 ... Hz</u> |
| **11. 2015** | 8.6 sec. segment<br>At O1, O2, Oz<br>0.53-150 Hz<br>1-sec epochs |  |  |  | **22.2%** B18 Cun<br>-25 -95 -5<br>**18.6%** B18 MOccG<br>20 -100 5<br>**9.2%** B19 MOccG<br>-50 -80 0 | <u>4.88, 9.77, 14.65, 20.51, 25.39, 30.27, 38.09, ... 59.57, 65.43, 69.34 ... Hz</u> |
| **15 2016** | 10.6 sec. segment<br>At O1, O2, Oz<br>0.53-70 Hz<br>1-sec epochs |  |  |  | **21.7%** B17 Ling<br>-10 -95 -20<br>**21.1%** B18 IOccG<br>25 -90 -15<br>**7.2%** B18 Ling<br>-5 -90 -20 | <u>7.81, 12.7, 15.63, 23.44, 30.27/31.25, 40.04, ... Hz</u> |
| **15 2018** | 8.3 sec segment<br>At O1, Oz, O2<br>0.53-150 Hz<br>1-sec epochs |  |  |  | **24.2%** B7 pCun<br>10 -80 40<br>**12.3%** B18 MOccG<br>-5 -100 10<br>**9.7%** B18 Ling<br>0 -80 5<br>**3.8%** B19 IOccG<br>45 -85 -10 | <u>4.88/5.86, 10.74, 16.1, 21.48, 26.37, 28.32... 60.55, 66.41... Hz</u> |
| **24. 2016** | 5.0 sec. segment<br>At O1, O2, Oz<br>0.53-70 Hz<br>1-sec epochs |  | **12.3%** B11 SFG<br>30 60 -10<br>**Slow wave cmpt** |  | **20.3%** B17 Cun<br>15 -90 5<br>**17.4%** B17 Ling<br>-10 -95 -20 | <u>6.84, 10.74, 13.67, 21.48, 28.32, 35.16, 41.99 Hz</u> |
| **26. 2016** | 50.0 sec. segment<br>At O1, O2, Oz<br>0.53-70 Hz<br>2-sec epochs |  | **9.4%** B47 IFG<br>50 45 -10<br>**Slow wave cmpt** |  | **29.7%** B18 Ling<br>-20 -100 -10<br>**10.9%** B17 Ling<br>5 -95 -5 | <u>3.42, 6.84/7.32, 10.74, 14.16, 17.58, 21.00/21.48 Hz</u> |
| **Sub-Totals**<br>Norm. to 100% |  | **7.9%**<br>**1.8%** | **35.2%**<br>**7.8%** | **7.9%**<br>**1.7%** | **399.0%**<br>**88.7%** |  |
| **ROIs**<br>Norm. to 100% |  | **B10 B11 B47 Frontal**<br>**9.6%** |  | **B6 Frontal near vertex**<br>**1.7%** | **B17 B18 B19 Occipital**<br>**88.7%** |  |

As far as this author is aware, there are no reported examples of harmonic structure in epileptic spike waves, which makes their observation in meditation all the more intriguing. Another major difference is the range of frequencies from 3.42 Hz to 7.81 Hz for meditation spike waves, significantly different to the 3-4 Hz range in absence epilepsy (Xanthopoulos, 2010) shown in the right-hand bar of Figure 6. And in source space the overwhelming occipital origin of meditation spike waves is strikingly different to the more widely varying locations, often frontal, in absence epilepsy (Stefan and Lopez da Silva, 2013).

#### 3.3.2. Seizure-Like or Epileptiform Activity

As noted in section 1.2.2, the transition from the first rūpa jhāna to the second and higher jhānas requires a meditator to become familiar with bodily energisation *pīti*. This typically develops in a natural way during samatha meditation, and we believe is evidenced by the high-energy ISWs we observe for some meditators. In the Yogāvacara tradition, however, specific techniques involving breath control are used to deliberately evoke such states for meditators with a special interest. The rationale is to become familiar with the ability to evoke higher energy states, and to then tranquilize that energy back into deeper absorption. To an observer, the subject typically displays clonic features similar to epilepsy, but without discomfort and able to evoke or leave the state at will. Figure 11 shows two examples from a wide range of presentations of this form of activity, analysis of which is still at an early stage. These examples are included to complete this overview and to illustrate broad features.

**Figure 11.**
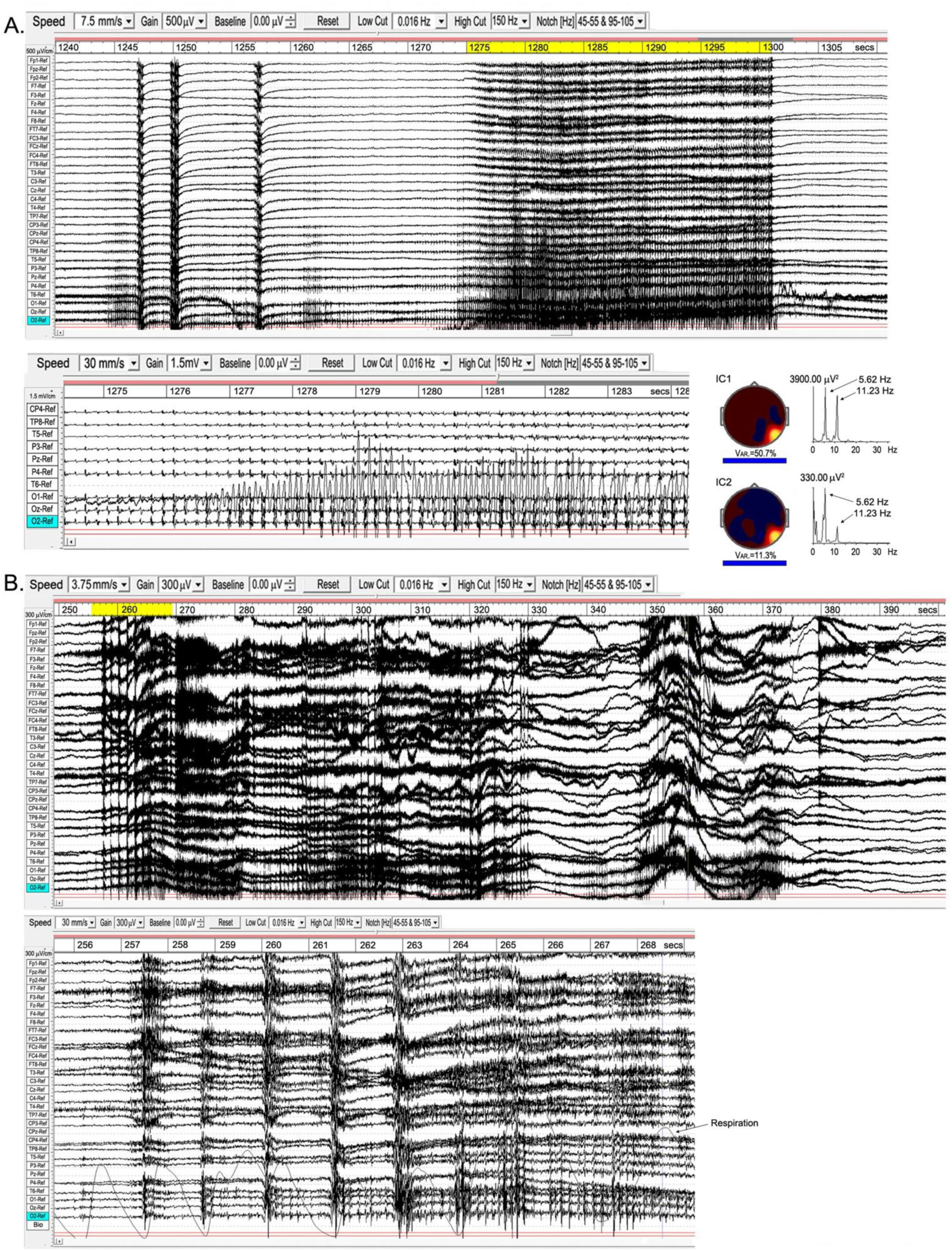
Examples of epileptiform activity. **A**, subject 15, 2018; the top panel shows the main 70-sec episode, with occipital activity expanded below; right is a source analysis of the 26-sec segment (yellow-highlighted) from the main episode. **B**, subject 7, 2015; the main 150-sec episode is above, with the early part of the “seizure” (yellow-highlighted) expanded below, together with respiration trace.

Example A is from an experienced practitioner. The first sign of increasing energisation is the development of occipital spike waves, followed immediately by a ∼½sec global ictal burst, a second burst 3 secs later, another ∼7 secs later, followed by the main body of the “seizure” 15 secs later. Physically, the meditator shows mild clonic jerks or bodily vibration, mainly along a vertical bodily axis. The expanded view below the top panel shows the occipital spike waves, with a related and near-sinusoidal rhythm at the right temporal site T6, reaching remarkable p-p values ∼3000 µV. To the right are the two strongest independent components (ICs) from an eLoreta analysis for the main 26-sec event, highlighted yellow. Activity is highly localised at Brodmann B37, MTG (MNI coordinates xyz 60 -60 - 5), with spike-wave frequency 5.62 Hz, with the temporal activity at the harmonic, 11.23 Hz.

Example B is more complex with stronger clonic activity. The EEG again shows brief global ictal bursts at onset, shown in the expanded view below together with respiration. Although these bursts coincide with the end of out-breaths, the regular breathing rhythm is not disturbed until the development of the paroxysmal effects are well-underway, and we do not believe hypercapnia/hypocapnia plays a role as it might in sleep disturbances. Occipital spike waves then develop progressively rather than precede the “seizure” as in example A. Probably because of the complex mix of activity in this example, including very strong ISW bursts as well as occipital spike waves and complex activity extending into the gamma band, a source analysis proved inconclusive, returning a range of small sources rather than any one or two dominant sources. Strong ISWs, as in this example, feature regularly in the more clonic examples of this activity, and we believe perform a containment function for the disturbance.

## 4. DISCUSSION

We have approached this series of case studies by focusing on specific paroxysmal EEG activity, not seen before, where the common factor is the practice of Buddhist jhāna meditation by subjects ranging from those with relatively early experience, to some with substantial experience. The detailed analysis of each of the observed themes in EEG activity will now be considered against the hypothesis that they relate to the nature of jhāna meditation, in a cross-discipline approach.

### 4.1. SPINDLES: EARLY DISRUPTION OF THE DCs, AND THE FIRST RŪPA JHĀNA

While meditation spindles have similar amplitudes to those in sleep, all other parameters are significantly different (Figure 2). Most strikingly, they are more prolific by an order of magnitude, and frequencies are substantially lower, towards the theta band. There is also evidence that more experienced meditators exhibit lower spindle frequencies. Cortical sources also differ: for meditation (Table 2) occipital sites are strong (23.2%), frontal sites weak (8.8%), with the remainder at fronto-parietal, limbic and temporal sites; while slow sleep spindles (10-12 Hz) are predominantly frontal, and fast sleep spindles (12-14 Hz; far higher than in meditation) largely parieto-temporal with smaller frontal and occipital contributions (Del Felice *et al*., 2014).

Spindling is also found in situations of conflicted attention (Sonnleitner *et al*., 2012), as well as during slow induction of propofol and volatile anaesthetics (Hagihira, 2017). In general anaesthesia frontal alpha increases, and under slow induction develops prolific spindling which fades as anaesthesia deepens (Hagihira, 2017; Hight, 2017; Gent, 2017). In all cases of anaesthesia a lowering of the alpha frequency is observed, with some evidence of longer spindle duration, as in meditation. However, spindles during slow induction are predominantly frontal, whilst in all our cases occipital sites are active, with varying degrees of extension over the scalp, but with frontal sites only weakly represented.

Sonnleitner *et al*’s (2012) study of driver-distraction and conflicted attention demonstrates enhanced alpha spindling even when drivers are not drowsy. In sensor space they find a broad distribution over the cortex, unlike in meditation. Since their study is relevant to driving, i.e. with eyes open, and subject to both visual and auditory distraction, it is not fully comparable to the eyes-closed meditation condition. No comment is made of any changes in frequency, and the descriptions are of “alpha spindles”.

All these modalities share a common theme of disruption to attention, either by driver distraction, chemically-induced anaesthesia, or the approach to sleep; or in the case of jhāna meditation the shift of attention away from our default sensory consciousness (DCs) as described in section 1.2.1. This strongly suggests that meditation spindles represent disruption to the cortical networks sustaining attention in the DCs, but in a manner significantly different to disruption in sleep or anaesthesia, which involve loss of consciousness. Also, given that spindling is by far the most common theme among our subjects (25 out of 27), we conclude that spindling is related to the early stages of developing jhāna that have to be negotiated *by all subjects* in order to access the higher jhānas. These are the development of access concentration and the first rūpa jhāna, representing growing success in resisting habitual attention processes of naming (inevitably related to language), recognition and discrimination (heavily visually-determined).

Since it is generally acknowledged that spindles represent thalamo-cortical interactions (Souza *et al*., 2016), we conclude that profound changes occur in related thalamo-cortical networks as meditators intentionally withdraw attention from their DCs. We might therefore expect involvement of networks related to the ventral and dorsal processing streams (Cloutman, 2012; Milner, 2017) heavily involved in attention and visual and auditory processing. The former carries rich and detailed representations of the world supporting cognitive processing; while the latter, sometimes referred to as the action stream, carries moment-to-moment information about objects and sense impressions as they relate to the experience of “I” or self. Enlarging on the discussion of section 1.2, it would fit rather well to link the aspect of attention described in jhāna texts as *vitakka* to the ventral stream, and the aspect of attention *vicāra*, which is much more feeling-based, to the dorsal stream. In addition to these two streams, we might also expect changes to networks supporting memory and spatial and temporal orientation, integral parts of our DCs experience of self. The spindle ROIs in source space summarised in Table 2 fit surprisingly well with this hypothesis, despite the limitations to spatial resolution of 31 or 21 electrodes. The ROIs are notably different to those in sleep (Del Felice *et al*., 2014), and the significant presence of limbic sources supports our expectation of effects on memory and spatial and temporal orientation, in accord with Kravitz *et al*’s (2011) view of limbic involvement in the ventral pathway. The main frequency range 7.5-10.5 Hz in meditation (Figure 2) corresponds well with the lower frequency ventral component of the alpha rhythm described by Barzegaran *et al*. (2017), which affirms our finding significant limbic sources underlying meditation spindles.

In our study we find no preference across subjects for either the dorsal or ventral paths in the balance of cortical sources, which may relate to this form of practice following a model of four rūpa jhānas (section 1.2.1) where the two aspects of attention, *vittaka* and *vicāra*, are worked on together in the first rūpa jhāna. An older and less well-known model (*Vimuttimagga*) describes five jhānas where *vitakka* is mastered in the first jhāna, *vicāra* in the second, with the third, fourth and fifth being the same as our second, third and fourth. In future recordings we may test this five-jhāna model. Even so, it may be relevant that in the underlying data supporting Figure 2, five subjects in the less-experienced cohort and six in the more experienced, show two peaks in the spectra of their EEG independent components, one in the alpha or low alpha band, and the other in the theta band; and we speculate that the alpha peak may relate to the activity of *vitakka* in disrupting the ventral stream, and the theta peak to the deeper activity of *vicāra* in disrupting the dorsal stream. In one of many resonances to work on active inference and predictive hierarchical processing in the brain, Kanai *et al*. (2015) describe two distinct processes, the first being a first-order driving process concerned with encoding the content of neuronal representations, and the second a second-order modulatory process that establishes context and salience. These two functions fit well if the former is related to the ventral stream and cognitive activities in *vitakka*, and the second to the dorsal stream and the feeling-based salience aspects of *vicāra*.

All the forms of attentional disruption described above have some relation to the alpha band. Meditation produces spindle activity from the lower alpha band down into the theta band; anaesthesia also lowers the alpha frequency; conflicted attention spindles remain within the alpha band, as do slow sleep spindles; whereas fast sleep spindles extend just above the alpha band. Foxe and Snyder (2011) review evidence that alpha activity may act as a sensory suppression mechanism in selective attention, while Jensen and Mazaheri (2010) see alpha activity performing an important gating mechanism in internetwork communication. Furthermore, Grandy *et al*. (2013) show that a person’s individual alpha frequency (IAF) is remarkably stable across periods of months, in response to a wide range of cognitive tasks. We therefore suggest that alpha activity, disrupted in all these modalities, is an integral part of the human DCs, far more important than simply an “idling rhythm”.

### 4.2. INFRASLOW WAVES (ISWs) AND THE HIGHER JHĀNAS

Meditation ISWs are significantly slower and higher amplitude than those in stage-4 nREM sleep (Figure 6), or even high-voltage delta coma (Libensen, 2012; Sutter and Kaplan, 2012). They also differ from slow delta activity in anaesthesia which is notably less rhythmic and coherent. In fact the highly developed rhythmicity and extensive nature of the strongest examples of meditation ISWs is very striking; particularly the tendency to form an annulus of alternating excitation-inhibition around relatively untouched central areas, apart from a localised region near the vertex. This surface pattern is quite different to the strong frontal predominance and smaller occipital focus in sleep (Bersagliere *et al*., 2017), and it is tempting to take a metaphor from sleep studies to infer that the annulus represents extensive areas of the cortex being “put to sleep”, or suppressed, by the high-voltage rhythmic ISWs; yet the subjective experience does not support this, since meditators report enhanced consciousness rather than any diminution.

In source space (Figure 8) we find a dominant midline vertex ROI, bridging the frontal-parietal divide, with frontal and occipital regions second and temporal sources third; very different to sleep, where Murphy *et al*. (2009) find the lateral sulci to be a major ROI, together with medial, middle and inferior frontal gyri, anterior and posterior cingulate, and precuneus; or Bersagliere *et al*’s (2017) eLoreta analysis similar to ours that finds predominantly frontal sources.

The mean amplitude 417±69 µV p-p for the strong examples of meditation ISWs is very significantly higher than other modalities, but in some cases can reach astonishing levels >2000 µV p-p, unprecedented as far as we are aware anywhere in the literature, for any modality, and representing a power ratio of four orders of magnitude increase from the resting state. We believe that such high amplitude ISWs reflect a deeper separation from the DCs than that characterised by spindling, corresponding in Buddhist understandings of jhāna to the increased energisation described by the jhāna factor *pīti*, heralding approach to the second rūpa jhāna, and subsequent development of the third and fourth jhānas. In this interpretation, those meditators in Table 1 showing both strong spindling *and* ISW activity would be developing experience of the first and second rūpa jhānas, while those subjects showing strong ISWs with either no or weak spindle activity would be consolidating their experience of the second rūpa jhāna, and those where powerful ISW activity dominates would be developing the second, third and fourth rūpa jhānas, culminating in the near absence of alpha activity and fully developed ISW activity near the vertex.

The experience of strong *piti* is always accompanied by heightened bodily sensitivity (section 1.2.2), and in some cases meditators report “waves of *piti*” moving across the head, which may relate to the slow travelling nature of these ISWs noted in the Results.

### 4.3 SPIKE WAVES

Whilst epileptic spike-wave bursts were formerly described as “generalised” rather than focal, often frontal, recent work such as Ji *et al.*’s (2015) fMRI study identifies specific thalamo-cortical networks having increased connectivity during such bursts. These include prefrontal-thalamic, motor/premotor-thalamic, parietal/occipital-thalamic, and temporal-thalamic networks. Similarly, Chiosa *et al*. (2017), using dense-array EEG, find significant connectivity changes between thalamus to frontal sites, and from frontal and temporal sites to thalamus preceding spike waves. However, meditation spike waves are overwhelmingly occipital, indicating a different mechanism, as does the range of frequencies (Figure 6) and harmonic structure, rather than the typical 3.0-4.0 Hz focal frequency of epileptic spike waves. As far as we are aware, there have been no reported findings of harmonic structure in the vast literature on epileptic spike waves, which makes our findings all the more intriguing.

The increased functional connectivity during epileptic discharges noted above is believed to reflect high levels of excitation and synchronization in one or more cortico-thalamic feedback loops, due to pathological conditions, locking the network(s) into fixed 3-4 Hz oscillation. In meditation, however, intentional rather than pathological, we suggest that meditation spike waves reflect a different form of destabilisation of the thalamo-cortical feedback loops when the personal element is withdrawn. And, further, because of the dominance of the occipital hub, that it is the “I/Eye” occipital-thalamic loop that is disrupted. The intentionally evoked “seizure-like” activity, including spike waves, presumably reflects the ability of some meditators to push the energisation (*pīti*) inherent to jhāna meditation beyond a threshold into instability. What is intriguing is that these phenomena do not appear to overly disturb a meditator’s tranquillity.

### 4.4 CONSCIOUSNESS

To date, research on the NCC has been limited to the human default sensory consciousness. This DCs is supported by a highly complex set of cortical networks in dynamic equilibrium to minimise free energy (Friston, 2010), with the personal component necessarily taking a central role. It should be no surprise that task-based studies reveal a sometimes bewildering array of NCC networks supporting the DCs.

We have noted the centrality of the alpha rhythm to spindling, and to the DCs, in effect as the *signature* of the DCs. Its ∼100ms periodicity is close to the human reaction time, with implied involvement of sensorimotor networks in readiness for action. In this sense the alpha rhythm performs the key time-synchronising role within DCs networks, which can be seen as biologically and evolutionary favoured for optimal response to threat and for species survival, providing a temporal scale factor for the DCs. The EEG frequency bands δ, θ, α, β and γ then become expressions of that scale factor, each independent and carrying different functions, but highly interconnected as postulated by Klimesch (2013). In terms of functions, it might be said that α activity characterises a minimum processing time for perception to action (a conscious “thought”); β the underling faster unconscious cognitive processes; θ the slower function of bridging more than one α process, facilitating memory; and γ representing higher-level functions perhaps related to consciousness of the whole.

The alpha scale factor may be considered an *attractor*, allowing otherwise scale-free neuronal activity to coalesce into DCs networks; we suggest that attentional disruption evidenced by spindles as meditators withdraw from the DCs reflects the beginnings of failure of the alpha attractor, leading to new network dynamics. As jhāna deepens with further withdrawal from the DCs, powerful ISWs develop and networks simplify further; first to near-midline frontal and occipital hubs with a developing vertex ROI, and as the higher jhānas develop to an eventually dominant near-vertex hub, with θ, α and β activity typical of the DCs virtually absent, leaving only broadband background gamma activity (Figure 9).

In the discussion on spindles, we mentioned the work of Kanai *et al*. (2015) who describe two distinct processes in predictive hierarchical processing, and we linked their first-order driving process that encodes the content of neuronal representations to *vitakka* in jhāna meditation, and their second-order modulatory process that establishes context and salience to *vicāra*. These two aspects of attention in the jhāna tradition would correspond well to the ventral and dorsal attention streams, which from this study appear to be the first disruptions to the DCs. We speculate that Kanai *et al*’s first-order process and the first jhāna factor *vitakka* relate to the outermost “shell” of the DCs; and that their second-order process that deals with salience and the first jhāna factor *vicāra* relate to the next higher-order part of the DCs to be disrupted; the effects of these disruptions being seen in the development of spindles. In our current four jhāna model we cannot yet separate these two effects in the EEG, although there is some supporting evidence in the two peaks observed for some meditators in the spectra of their independent components.

As attention becomes stabilised and the second jhāna develops, cortical activity simplifies into a frontal-occipital axis with a smaller temporal contribution, with a progressively dominating ROI near the vertex (Figure 8). These cortical hubs are interesting when considered against the nature of jhāna as progressively moving from a sensorily-determined subject-object basis of consciousness, towards the inner absorption common to all the jhānas (sections 1.2.2, 1.2.3). On that basis, we suggest that the occipital and frontal hubs represent the residual subject and object poles, respectively, of the sensory DCs, the temporal contribution the residual part of the ventral perceptual stream, while the vertex hub is a sign of the emerging second rūpa jhāna consciousness. In this model, the occipital hub, integral to the dorsal and ventral perceptual streams of the DCs, carries the first-person “I/eye” pole of sensory consciousness (Merker 2013), while the frontal hub and associated executive attention network carries the object pole due to its role in cognitive processing (Petersen and Posner, 2012), the two relating to each other through recurrent connectivity. The occurrence of spike waves for some meditators was described earlier as reflecting disruption to the “I/Eye” occipital-thalamic, which lends support to this.

As a meditator progresses towards the higher jhānas, the near-vertex hub eventually dominates, encompassing Brodmann sites, B6, B5 and B7. These include the supplementary motor area (SMA), strongly connected to the thalamus and projecting directly to the spinal cord, and the highly connected medial parietal associative cortex, believed to be widely involved in high-level processing tasks, with dense links to the underlying cingulate, thalamus and brain stem. We might also include the role of the anterior cingulate cortex in predictive coding discussed in Seth *et al*. (2012) in relating *presence* to agency, with resonance to meditators’ descriptions of “vivid presence” during the deeper stages of jhāna. This vertical-axis connectivity is striking in contrast to the (suggested) front-back axis of the DCs, and suggests involvement of the ascending reticular activating system (ARAS), with its known involvement in arousal, attention and consciousness (Maldonato, 2014), and again likely related to the processes of top-down–bottom-up recurrent processes in active inference (Friston *et al*., 2016; Seth *et al*., 2012; Seth and Friston, 2016). It is known that disruptions to the ARAS can lead to coma (Norton, 2012), and disruption of posterior cingulate connectivity can cause unconsciousness (Herbet, 2014), but the form of disruption we see in this form of meditation does not lead to unconsciousness, even though the strong ISWs have superficial similarities to deep sleep and coma, as discussed earlier.

As noted in section 1.2.3, the third rūpa jhāna is described as “completely conscious” in the 5^th^-century *Vimuttimagga*, which quality continues into the fourth rūpa jhāna. We believe the fully developed near-vertex hub of powerful ISWs reflects jhāna consciousness with DCs lower frequency activity now virtually absent (Figure 9). From our study, jhāna consciousness appears to be characterised by the scale factor of the respiration cycle, not entirely a surprise given that the breath is the primary object in the early stages. However, as a scale factor this is almost two orders of magnitude slower than the 100 ms of the DCs, which might explain the expanded sense of time and spaciousness meditators describe during this form of meditation, and the frequent rendering of Samatha as undisturbed peace or tranquillity. The development of a vertex-body axis, rather than the frontal-occipital axis of the DCs, and evidence of an even slower underlying ISW activity ∼0.02 Hz, suggests a slow metabolic scale factor integrating the entire mind-body system in the deeper levels of jhāna. This very slow activity may correspond to the slow alternations seen in fMRI*/*BOLD imagery of subjects in the resting state (Grooms *et al*., 2017), and may represent a harmonic “beating” between other ISW frequencies present, similar to Steyn-Ross *et al*.’s (2011) model for ultra-slow oscillations.

The sense of *presence* reported by meditators is an embodied presence, resonating with the phrase *embodied selfhood* in Seth and Friston’s (2016) work on active interoceptive inference. The intense vertex focus, unlike the dual frontal-occipital ROIs of the DCs, raises the interesting question as to what is the nature of the subject-object structure of jhāna consciousness, since an object is still required to be *conscious of*, and Buddhist texts are quite clear in regarding the jhānas as states of consciousness. In the oral jhāna tradition several views are expressed. One is that each moment of consciousness becomes the object of the next, giving the illusion of perfectly still and continuous undisturbed consciousness. This is envisaged as a high level and very fast process, and we might wonder at the role of the background gamma activity and brief gamma bursts we observe. We might also consider that it might represent a highly stable state of reciprocal top-down–bottom-up recurrent processes (Friston *et al*., 2016) where the error between prediction and current state has been reduced to effectively zero, at least for a while. A second view is that the body itself is the supporting object of jhāna consciousness, as part of a deep metabolic integration, and perhaps related to the intriguing term “body witness” encountered in the ancient texts (e.g. *Vimuttimagga*).

#### 4.4.1 Scale factors and harmonic structure

Whilst the scale factor for the DCs is suggested to be the ∼10 Hz alpha rhythm, that for SWs in sleep or coma is determined by haemodynamic pressure (Mensen *et al*., 2016), where the scale factor is the heart rhythm, with corresponding SW frequencies ∼1.0-1.2 Hz similar to typical pulse rates. Meditation ISWs, however, appear to be related to the respiration rhythm, with mean frequencies ∼0.125Hz.

Meditation spike waves show harmonic spectral structure, unlike those in epileptic or pathological states, and we have suggested that their appearance relates to withdrawal of the personal “I” component from the occipito-thalamic network. We further suggest that this disruption triggers the thalamus into harmonic activity in an attempt to stimulate scaled network activity similar to that of the DCs; the implication being that harmonic or fractal structure is an integral part of thalamo-cortical network connectivity, and that the δ, θ, α, β structure of the DCs is but one example, that we presume is optimal for sensory consciousness and minimisation of free energy.

### 4.5. MODERN NEUROSCIENCE, ANCIENT JHĀNA

The results and preceding discussion speak to a close relationship between the different aspects of consciousness that reflect departures from our default consciousness, and their neurophysiological correlates, in the light of understandings of the ancient practice of jhāna meditation. Furthermore, these correlates highlight the key role of attention and concomitant changes in neurophysiological excitability. This is entirely consistent with modern-day formulations of hierarchal inference in the brain; particularly in terms of internal attention states, interoceptive inference and the relationship between consciousness and sleep. The common theme is a physiological modulation of cortical excitability and ensuing neuronal dynamics that under the free-energy principle and Bayesian brain hypotheses play the role of precision weighting (Clark, 2013; Hohwy, 2013; Seth and Friston, 2016). In brief, precision weighting acts to date newsworthy signals (e.g., ascending prediction errors in cortical hierarchies) according to their reliability or salience. Psychologically, this has often been construed in terms of both exogenous and endogenous attention (Chawla et al., 1999; Feldman and Friston, 2010) and, on some readings, the essential neurophysiology of consciousness itself – and a sense of selfhood (Palmer et al., 2015; Seth, 2014; Seth et al., 2012; Stephan et al., 2016). Interestingly, the neuromodulatory control of excitability (and implicitly precision) is thought to underwrite the shifts in conscious states during the sleep-wake cycle, leading to a tight relationship between consciousness, sleep and dreaming (Hobson and Friston, 2012 and 2014). From a purely computational perspective, mindfulness, meditation and (possibly) sleep may be seen as representing brain states partially divorced from the sensorium (by assigning sensory signals very low precision), or, in the language of this paper, partially separated from the DCs.

The importance of these brain states is to enable minimisation of free energy via the minimisation of the statistical construct called complexity in the active inference literature. In essence, this simply means that redundant connections can be removed from the brain’s hierarchical generative models; thereby enabling the brain to generalise its models or explanations for exchange with the world – when re-engaged by sensory input. On this view, one could regard this form of meditation as aspiring to that simple but serene state where all that is redundant, distracting or unnecessarily complex can be dissolved. In practice, we find that the progressive and extensive simplification of network activity as a subject develops jhāna, leads to highly focused activity near the vertex, suggesting extensive connectivity down through the brain stem into the body, reflecting the subjective experience of embodied presence, and the meditational terms *samādhi* or yoga that describe a deeply integrated mind-body experience. It is as though all the previously widely distributed networks, each consuming portions of available energy, are gathered together into a simple focus revealing a previously unrecognised and remarkable reservoir of available energy.

We believe that the capacity to manage and contain such high-energy states is the result of the detailed development of attention, often over many years, coupled with the deliberate use of non-normal lengths of breath as noted in the Introduction. Indeed, we wonder whether some of the basic features of this attentional development might be adapted for epilepsy sufferers with the goal of reducing frequency of seizures. The study also demonstrates an equally remarkable *responsiveness* of the brain’s cortical networks to willed intensions of subjects during this form of meditation, which we presume results from the temporary experience of greatly reduced complexity noted above, and is no doubt also related to plasticity.

## ACKNOWLEDGEMENTS

The author is grateful for the generous cooperation of all the subjects who participated in this study, for the ethical approval granted by The Samatha Trust, and to Stefano Caria for sharing his expertise and advice on statistical methods.

